# Purinergic Receptor P2Y_13_ Controls Activation and Mode of Division in Subependymal Adult Neural Stem Cells

**DOI:** 10.1101/2024.11.29.626065

**Authors:** Lucía Paniagua-Herranz, Julia Serrano-López, Celia Llorente-Sáez, Marina Leonor Pérez-Sanz, David de Agustín-Durán, Pere Duart-Abadia, Ana Domingo-Muelas, África Vincelle-Nieto, Paloma Bragado, Diana Manzano-Franco, Marina Arribas-Blázquez, Lucía Gallego, Luis Alcides Olivos-Oré, Sergio Gascón, Álvaro Gutierrez-Uzquiza, Raquel Pérez-Sen, Esmerilda G. Delicado, Rosa Gómez-Villafuertes, Armando Reyes-Palomares, Antonio R. Artalejo, Isabel Fariñas, Felipe Ortega

## Abstract

The subependymal zone (SEZ) of the mammalian brain is the most active germinal area that continues to generate newborn neurons throughout life. This area harbors a population of neural stem cells (NSCs) that can be found in different states of activation, each differing in proliferative capacity and molecular signature: quiescent NSCs (qNSCs), primed NSCs (pNSCs), and activated NSCs (aNSCs). There is currently a void in terms of the specific markers available to effectively discern between these transient states. Likewise, the molecular signaling mechanisms controlling the transition from quiescence to activation remain largely unexplored, as do the factors influencing the decision between differentiation and self-renewal during NSC division. Here, we present evidence that the metabotropic P2Y_13_ purinergic receptor plays a critical role in regulating adult neurogenesis. We found that P2Y_13_ is specifically expressed in NSCs within the adult SEZ and that its levels can be used to distinguish qNSCs from aNSCs. Functionally, P2Y_13_ signaling promotes NSC activation, enhancing lineage progression, while dampening their self-renewal capacity. Conversely, pharmacological blockade or genetic silencing of the P2Y_13_ receptor favors NSC quiescence. Thus, we identified the metabotropic P2Y_13_ purinergic receptor as a pivotal modulator of NSC dynamics, influencing both the balance between NSC quiescence and activation and the mode of NSC division at the subependymal zone.

## INTRODUCTION

The generation of newborn neurons within the central nervous system (CNS) is a process not limited to embryonic development but rather, it continues in the mammalian brain throughout life. Neurogenesis takes place in specialized microenvironments called neurogenic niches(Obernier & Alvarez-Buylla, 2019), of which two stand out in the adult: the subgranular zone in the hippocampal dentate gyrus (DG), and the subependymal zone (SEZ) located in the walls of the lateral ventricles(Lim & Alvarez-Buylla, 2014). The SEZ is the largest and most active germinal region, harboring the vast majority of CNS neural stem cells (NSCs). These cells are derived from embryonic radial glia that enter quiescence at midgestation until their activation in the adult (Fuentealba *et al*, 2015; Furutachi *et al*, 2015) and they are characterized by astroglial features, such as expression of the glutamate-aspartate transporter (GLAST) and glial-fibrillary acidic protein (GFAP). When activated, subependymal NSCs give rise to transit amplifying progenitor cells (TAPs), which divide three or four times before giving rise to neuroblasts (NBs). These NBs are considered to be immature neurons and they subsequently migrate through the rostral migratory stream (RMS) to become mostly adult-generated GABAergic interneurons in the olfactory bulb (OB) (Costa *et al*, 2011; Obernier & Alvarez-Buylla, 2019; Ponti *et al*, 2013).

Beyond the mosaic organization of the neurogenic niche (Merkle *et al*, 2014; Merkle *et al*, 2007), the intricate complexity of the SEZ cytoarchitecture has become more evident as the heterogeneity and mode of division of the NSC pool has been revealed. NSCs can be found in three molecularly distinct states: quiescent NSCs (qNSCs), NSCs that are still quiescent but that are “primed” for proliferation (pNSCs), both EGFR^low/-^ and with a quiescence-associated transcriptome, and activated EGFR^+^ NSCs (aNSCs) (Belenguer *et al*, 2021b; Chaker *et al*, 2016; Dulken *et al*, 2017; Llorens-Bobadilla *et al*, 2015; Mizrak *et al*, 2019). Despite recent efforts to generate reliable protocols to isolate subependymal NSCs in different states (Basak *et al*, 2018; Belenguer *et al*, 2021a; Cebrian-Silla *et al*, 2021; Codega *et al*, 2014; Dulken *et al*., 2017; Giachino *et al*, 2014b; Llorens-Bobadilla *et al*., 2015; Mizrak *et al*., 2019), the lack of specific markers prevents the individual contribution of each type of NSC to neurogenesis from being defined, also impeding an assessment of their maintenance over time. Furthermore, quiescence-activation dynamics and the mode of division following activation still remain incompletely understood. GFAP-based lineage tracing has indicated that NSC activation results in asymmetric division (Calzolari et al., 2015) whereas a more recent analysis reports that NSCs undergo consuming differentiative divisions 80% of the times, while only 20% of aNSC divisions are self-renewing (Obernier et al., 2018). Both, the transition from quiescence to activation and the mode of division are critical in modulating the neurogenic potential of the SEZ niche. These features could be targeted to reactivate the niche during aging or enhance the therapeutic potential of these cells for brain repair in pathological scenarios. However, the mechanisms by which signalling pathways regulate these NSC traits and their coordination remain poorly understood (Obernier & Alvarez-Buylla, 2019).

Quiescent NSCs express receptors for multiple signals but only a few have been shown to activate NSCs out of their quiescent state (Blasco-Chamarro & Farinas, 2023; Doze & Perez, 2012; Kobayashi & Kageyama, 2021; Urban & Guillemot, 2014). Purinergic signaling is one of the oldest cell-to-cell communication systems known, driven by extracellular nucleotides (mainly ATP) and their interactions with specific cell-surface purinoceptors (P2 receptors). Although adenine nucleotides [ATP, adenosine diphosphate (ADP), and adenosine (ADO)] were initially described as neurotransmitters in neural circuits, they function in a variety of cells as signaling molecules. Purinergic receptors are divided into P2X receptors, ligand-gated calcium channels and P2Y receptors, the latter a sub-family of eight G-protein coupled metabotropic receptors in mammals (P2Y_1,2,4,6,11,12,13,14_) (Abbracchio *et al*, 2006; Burnstock, 1972; Burnstock *et al*, 1970). The balance between the release of extracellular nucleotides, mostly through the action of the vesicular nucleotide transporter (VNUT)(Miras-Portugal *et al*, 2019; Sawada *et al*, 2008), their interaction with the purinergic receptors in each cell population and their degradation by extracellular nucleotidases, defines the role exerted by this signaling pathway. Within the CNS, the purinergic system is involved directly in development and in neurogenic niches in particular (Oliveira *et al*, 2016; Ortega *et al*, 2021; Paniagua-Herranz *et al*, 2020b; Zimmermann, 2006). In fact, elements of the purinergic system are known to regulate NSC maintenance (Weissman *et al*, 2004) and indeed, the P2X7 purinergic receptor co-localizes with the radial glia palisades present in the brain circumventricular organs during mouse embryonic development (Ortega *et al*., 2021). Interestingly, we also confirmed a strong influence of the purinergic system in post-natal cerebellar neurogenic niches, where it is involved in controlling both neurogenic activation and self-renewal within the NSC population. Importantly, purinergic signaling was effective before the onset of canonical neurotransmitter expression. Indeed, maximal VNUT expression correlated with the peak of post-natal neurogenesis, and it was present in the neurogenic niche before vesicular glutamate or GABA transporter expression was detected in the developing cerebellum (Paniagua-Herranz *et al*., 2020b). Thus, NSCs may preferentially use purinergic communication, even before other neurotransmitter systems are fully established during development. Consequently, the purinergic system postulates itself as a primary cell-to-cell communication network within neurogenic niches that allows NSC lineage progression to be fine-tuned.

Here we show that the metabotropic P2Y_13_ purinergic receptor plays a critical and specific role in regulating adult neurogenesis. We found that P2Y_13_ is specifically expressed by the NSC population in the adult SEZ, and that its expression can differentiate qNSCs from aNSCs. At the functional level, signaling through P2Y_13_ promotes NSC activation enhancing lineage progression and neurogenesis while reducing NSC self-renewal capacity. Conversely, pharmacological blockade or genetic silencing of the P2Y_13_ receptor promotes NSC quiescence. Altogether, our results indicate that the P2Y_13_ receptor is a key element in the decision of an NSC to undergo differentiative divisions when it activates. P2Y_13_ receptors, therefore, regulate the balance between quiescence and activation of qNSCs in coordination with the modulation of NSC division mode in the SEZ.

## RESULTS

### P2Y_13_ levels distinguish quiescent from activated NSCs in the adult SEZ

P2Y purinergic receptors (P2Y_1,2,4,6,11,12,13,14_) activated by extracellular nucleotides can be divided according to their preferential ligands and the signaling pathways they couple to (Jacobson *et al*, 2020). While the P2Y_12_ receptor is preferentially expressed by microglia and vascular cells (Prinz *et al*, 2019; Rauch *et al*, 2010; Simon *et al*, 2002; Wihlborg *et al*, 2004), P2Y_1_ has been implicated in cortical development (Weissman *et al*., 2004) and P2Y_13_ in the modulation of adult neurogenesis in the DG, as well as in neuronal differentiation and survival (del Puerto *et al*, 2012; Espada *et al*, 2010; Morente *et al*, 2014; Ortega *et al*, 2011b; Perez-Sen *et al*, 2017; Perez-Sen *et al*, 2015; Stefani *et al*, 2018). RT-PCR analysis of subependymal homogenates indicated that P2Y_13_ receptor mRNA was the most strongly expressed of the metabotropic receptors (**Figure 1A**). P2Y_13_ receptor expression was further examined in the dorsal and ventral walls of the SEZ independently, two areas with distinct potentiality, such that the dorsal walls are prone to oligodendrogliogenesis while neurogenesis is promoted in the ventral walls (Colak *et al*, 2008; Menn *et al*, 2006; Ortega *et al*, 2013b). Although no differences in P2Y_13_ transcript expression was evident between these areas, significantly more P2Y_13_ protein accumulated in the ventral wall (**Figure 1B, C**), suggesting its possible involvement in neurogenic lineage progression. As a result, the expression of the P2Y_13_ receptor within the neurogenic lineage was analyzed using GFAP as a marker of mature astrocytes and NSCs, the Achaete-Scute Family bHLH Transcription Factor 1 (Ascl1) to label TAPs, and βIII-tubulin to label NBs (Obernier & Alvarez-Buylla, 2019). Immunohistochemical (IHC) analysis of the ventral wall of the SEZ revealed that P2Y_13_ co-localized exclusively with GFAP^+^ astroglia, and it was not detected in either NBs or TAPs (**Figure 1D-F**). The same pattern was also observed when NSCs and their progeny were analyzed in culture, acutely isolated from their niche (**Figure S1**). Moreover, we confirmed that the P2Y_13_ protein precisely co-localized with GFAP^+^/SOX2^+^ NSCs (**Figure 1G**). To assess for the presence of this receptor in the different states of NSC activation, we used a modified version of the previously published flow cytometry strategy to fractionate the entire neurogenic lineage (Belenguer et al., 2021a; b). Since P2Y_13_ is strongly affected by enzymes, we dissociated the tissue solely by mechanical dissociation. Although this procedure significantly increased cell death, we could clearly distinguish a significantly stronger presence of P2Y_13_ in the quiescent EGFR^low/-^ NSC population than in the EGFR^+^ aNSC population, again with no labeling of NBs (**Figure 1H**; **Figure S2**). Together, these data suggest that the P2Y_13_ receptor may potentially serve as a biomarker to specifically label and isolate not only NSCs but more specifically, the qNSCs within this population.

**Figure 1.**
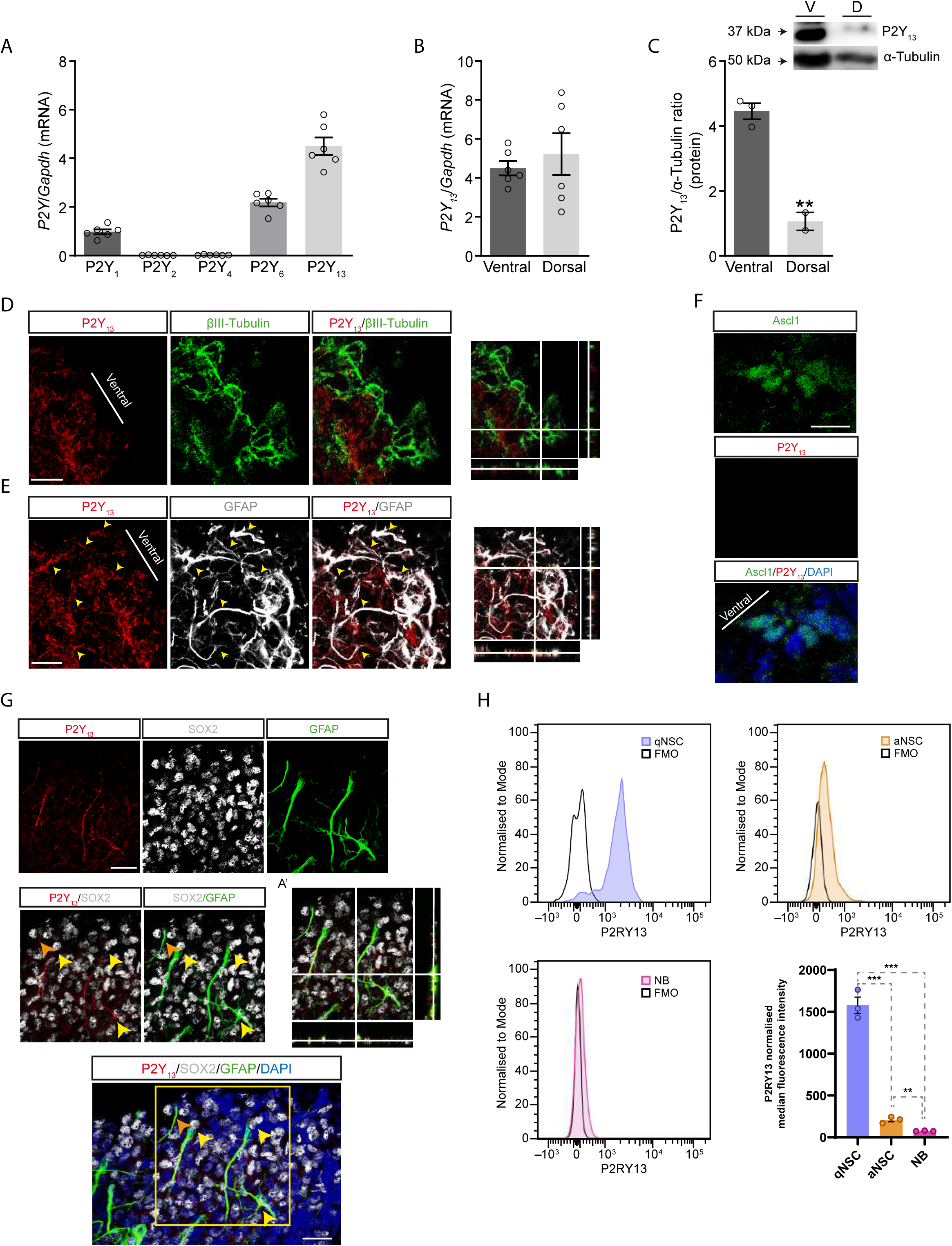
The P2Y_13_ receptor is expressed by qNSCs in the adult SEZ. **A.** The expression of the different metabotropic P2Y receptors in the adult SEZ was analyzed by quantitative RT-PCR (n=6). **B.** P2Y_13_ receptor expression in the ventral and dorsal wall of the adult SEZ analyzed by quantitative RT-PCR (n=6). **C**. Comparison of the P2Y_13_ receptor protein in the ventral and dorsal wall of the SEZ (n=3). **D-F**. P2Y_13_ receptor expression in the ventral wall of the SEZ. Note how the P2Y_13_ receptor (red) co-localizes with GFAP positive (white) astroglia (yellow arrowheads) but not with Ascl1 positive TAPs or cells expressing βIII-tubulin (green, scale bar 30 µm except for Ascl1 10 µm).**G**. Co-localization of the P2Y_13_ receptor (red) in NSCs with SOX2 (white) and GFAP (green) in the adult SEZ (scale bar 30 µm). **H.** SEZ-derived cell populations expressing the P2Y_13_ receptor in a FACs analysis using a P2Y_13_-GFP conjugated antibody. Note how the expression is mainly associated with the qNSC fraction (n=3). All graphs show the mean ±SEM: *p<0.05, **p<0.01, and ***p<0.001 (T-test).

### P2Y_13_ is functional on SEZ-derived NSCs

P2Y receptors are known to promote calcium release from the endoplasmic reticulum into the cytoplasm (Abbracchio *et al*., 2006). Calcium microfluorimetry experiments were, therefore, performed to assess whether the P2Y_13_ receptor was fully functional in adult NSCs. To do so, we took advantage of a method developed in house that enables the cell culture of NSCs and their progeny, isolated from their neurogenic niche signals and in the absence of added mitogens. Importantly, isolated NSCs maintain their neurogenic potential despite the absence of signals from their niche turning this approach into a unique tool to assay the effects of specific signaling molecules as P2Y_13_ agonists (Costa et al., 2011; Gomez-Villafuertes et al., 2017; Ortega et al., 2013a; Ortega and Costa, 2016; Ortega et al., 2011a). Exposure to the selective P2Y_13_ agonist 2MeSADP (10 µM) induced an increase in intracellular calcium in cultured NSCs, an effect that was sensitive to the presence of the specific antagonist of P2Y_13_ receptors MRS2211 (10 µM), but not to MRS2179 (10 µM), a specific antagonist of the P2Y_1_ receptor that can also be activated by 2MeSADP (**Figure 2A, B**). Remarkably, post-imaging immunocytochemistry (ICC) in the same cells confirmed that only GFAP^+^/SOX2^+^ NSCs reacted to the agonist, while parenchymal astrocytes expressing only GFAP and cells expressing only SOX2 (presumably TAPs or early NBs) were not affected by the presence of 2MeSADP (**Figure 2C**). Further evidence of the specificity of this response was obtained by analyzing the downstream signaling triggered by P2Y receptors, which can modulate the outward currents of potassium channels (Coppi *et al*, 2012; Shrestha *et al*, 2010; Wu *et al*, 2012). This feature was assessed in conjunction with an inherent hallmark of NSCs in culture, their marked cell growth prior to division that exceeds that of other neurogenic lineages (Costa *et al*., 2011). Accordingly, in patch-clamp recordings obtained from the cells with the highest growth rates over 24 h in SEZ cell cultures, the outward currents developed by these cells were potentiated by 2MeSADP, an effect that was impeded in the presence of MRS2211 but not MRS2179 (**Figure 2D, E**). These data indicate that under non mitogenic conditions, cultured NSCs have functional P2Y_13_ receptors.

**Figure 2.**
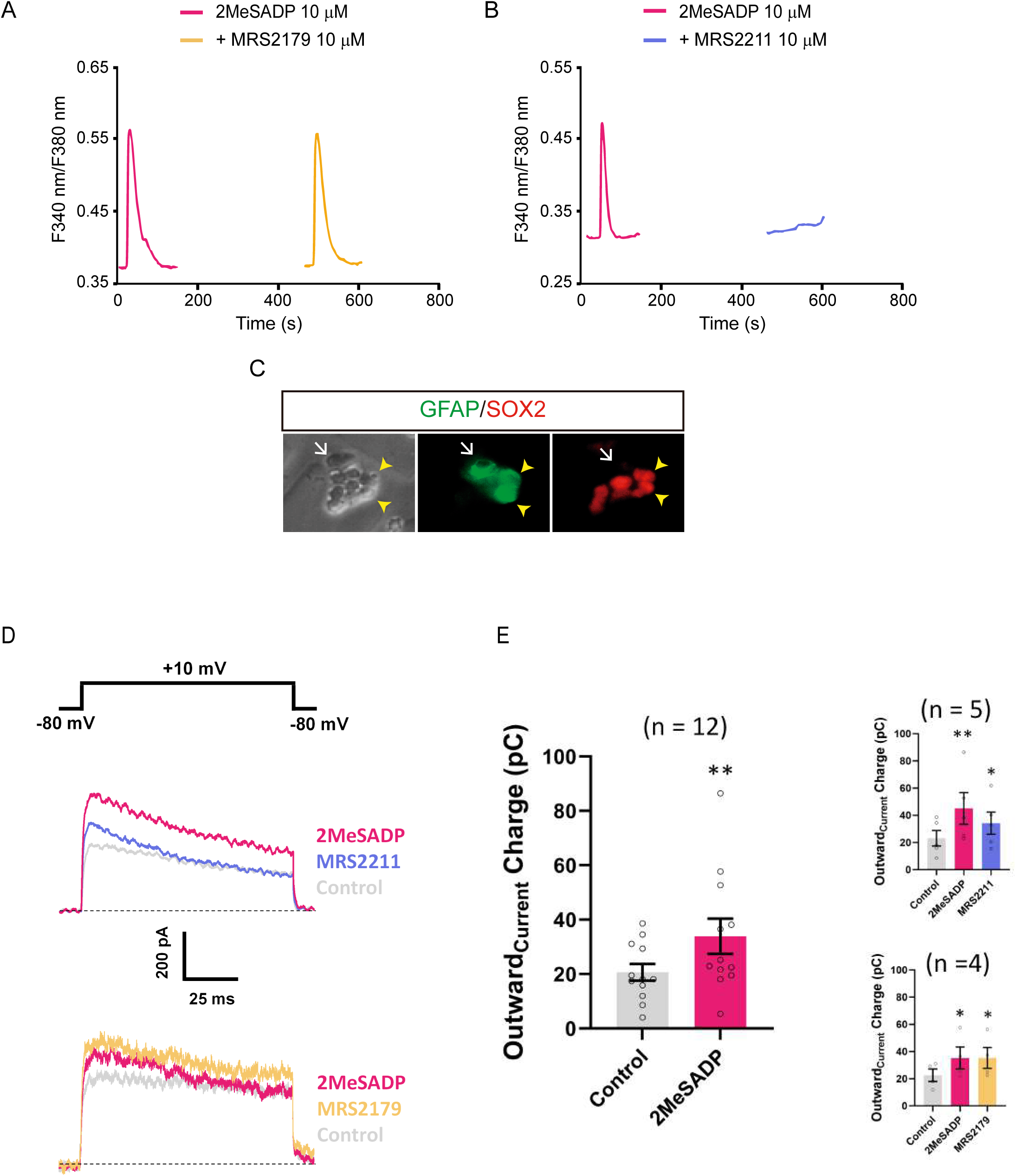
The P2Y_13_ receptor is active in SEZ-derived NSCs in culture. NSCs in culture and loaded with the calcium dye Fura-2 were stimulated with 2MeSADP and after a washout period, they were re-exposed to 2MeSADP in the presence of either the P2Y1 receptor antagonist MRS2179 **A** or the P2Y_13_ receptor antagonist MRS2179 **B**. All compounds were tested at a concentration of 10 μM and representative traces of the F340/F380 fluorescence ratios recorded from single cells are shown. **C**. Immunocytochemistry identifying GFAP (green) and SOX2 (red) double positive NSCs that respond to the selective agonist of the P2Y_13_ receptor, 2MeSADP (yellow arrowheads). Conversely, the intracellular calcium levels of GFAP positive parenchymal astrocytes (white arrow) does not change. **D**. Effects of 2MeSADP on voltage-gated currents in SEZ-derived NSCs in culture. Outward currents evoked by a depolarising pulse (+10 mV, 100 ms from a Vh of −80 mV) were increased in the presence of 2MeSADP (10 µM, 2 min). Subsequent exposure to MRS221 (10 µM, 2 min) partially reversed the potentiating effect of 2MeSADP (upper panel). At variance, exposure to MRS2179 (10 µM, 2 min) of 2MeSADP with MRS2179 (10 µM, 2 min) did not have any effect on outward current increase elicited by 2MeSADP (10 µM, 2 min) (lower panel). **E**. Scatter plot of outward current charges from the experiments shown in (**D**), in the presence or absence of 2MeSADP (10 µM, 2 min: left panel), and following co-incubation (right panels) with MRS2211 (upper right) or MRS2179 (lower right). The values are the means ± SEM of the number of cells indicated between parentheses; the statistical significance was assessed using the student’s T-test for paired samples: *p < 0.05; **p < 0.01.

### P2Y_13_ modulates NSC behavior *in vivo*

Because the release of nucleotides is tonic (Lin *et al*, 2007; Young *et al*, 2011), we next addressed whether P2Y_13_ activity also influences NSC behavior and lineage progression in the adult SEZ *in vivo* by inactivating P2Y_13_ expression. Moreover, the specific expression of this receptor in qNSCs prompted the question as to whether this receptor might be involved in establishing the boundaries between quiescence/activation and subsequent neurogenic progression. Thus, in order to uncover the physiological role of P2Y_13_ receptors, CRISPR/Cas9 lentiviral constructs were designed (sgP2Y_13_) to stably delete their expression and dampen their activity in the adult SEZ (**Figure S3**). Control (gNTC) or sgP2ry_13_ lentiviruses, which target proliferating as well as non-proliferating cells, were injected into the ventral wall of the adult SEZ together with lenti-Cas9 lentiviruses. IHC staining to visualize Cas9+ cells 14 days post-injection (dpi) revealed a significant increase in the percentage of GFAP^+^/SOX2^+^ cells within the sgP2ry_13_/Cas9 positive cell population in the ventral wall of the SEZ (28.71 ± 2.28% in sgP2Y_13_ and 11.29 ± 2.53% in gNTC; **Figure 3A, B**), suggesting that P2Y_13_ activity may influence the activation of NSCs. Quantification of Cas9+/Ki67+ cells with respect to the total number of Cas9 positive cells confirmed that this increase was not due to enhanced proliferation of the sgP2ry_13_/Cas9 population (**Figure 3C, D**). Moreover, the increase in NSCs was not coupled to an increased in the number of Cas9^+^/DCX^+^ neuroblasts (8.04 ± 0.55% gNTC, 5.23 ± 1.35% sgP2Y_13_; **Figure 3E, F**). These results suggest that the silencing of the P2Y_13_ receptor may instruct the NSCs to remain quiescent within the adult SEZ.

**Figure 3.**
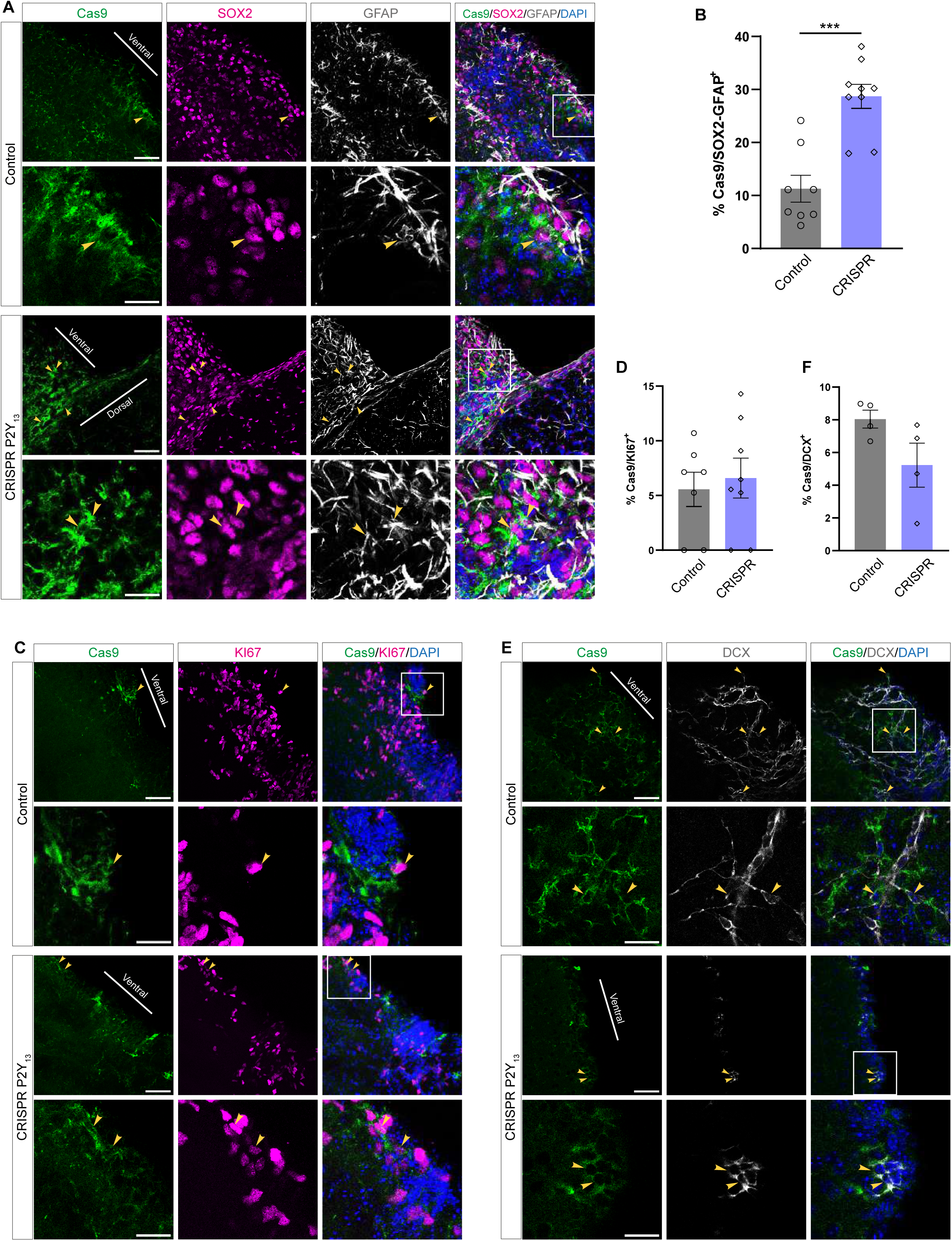
P2Y_13_ silencing increases the number of NSCs remaining in the SEZ without promoting proliferation. **A.** The effect of local P2Y_13_ receptor silencing on the NSC population, achieved using the CRISPR/Cas9 sgP2Y_13_ lentiviral vectors: Cas9 (Green), SOX2 (Magenta) and GFAP (White), the cell nuclei are stained with DAPI. The lower panels show a higher magnification of the selected areas (scale bar 50 µm). **B.** Quantification of the Cas9^+^/GFAP^+^/SOX2^+^ cells remaining in the adult SEZ (n=4). **C** Effect of the local silencing of the P2Y_13_ receptor using the CRISPR/Cas9 sgP2Y_13_ lentiviral vectors on the proliferation within the SEZ: Cas9 (Green), Ki67 (Magenta) and GFAP (White), the cell nuclei are stained with DAPI. The lower panels show a higher magnification of the selected areas (scale bar 50 µm). **D**. Quantification of the Cas9^+^/Ki67^+^ cells SEZ (n=4). **E**. Effect of local silencing of the P2Y_13_ receptor using the CRISPR/Cas9 sgP2Y_13_ lentiviral vectors on the neuroblast population: Cas9 (Green), DCX (White) and the cell nuclei are stained with DAPI. The lower panels show the higher magnification of the selected areas (scale bar 50 µm). **F.** Quantification of the Cas9^+^/DCX^+^ cells (n=4). All graphs show mean ±SEM: ***p<0.001 (T-test).

To further address this point we performed gain-of-function experiments with lentiviruses carrying a bicistronic construct containing the P2Y_13_ receptor cDNA and a GFP reporter (LV-P2Y_13_-iresGFP; **Figure S4A**) to achieve local expression of the P2Y_13_ receptor. Injection of LV-P2Y_13_-iresGFP into the ventral wall of the adult SEZ was confirmed to induce overexpression of these receptors in GFP^+^ cells, unlike lentiviruses encoding GFP alone (LV-GFP; **Figure S4B, C**). As commented before, it is important to note that a more prominent presence of the receptor in NSCs will be coupled to stronger P2Y_13_ activation due to the tonic release of nucleotides within the adult SEZ (Lin *et al*., 2007; Young *et al*., 2011). Thus, the relative distribution of the GFP^+^ positive cells in the ventral wall of the SEZ and in the RMS 14 days after injection of LV-P2Y_13_-iresGFP was examined as a readout of the activation/differentiation balance among the transduced cells. Lentiviral expression of the P2Y_13_ receptor significantly enhanced the relative proportion of GFP^+^ cells in the RMS relative to that driven by LV-GFP, consistent with enhanced activation and NB lineage progression in the SEZ (**Figure 4A-C**). The number of GFP^+^/GFAP^+^/SOX2^+^ NSCs remaining in the ventral wall of the SEZ 14 days after injection was significantly decreased after LV-P2Y_13_-GFP injection (**Figure 4D-F**), supporting the notion that the P2Y_13_ receptor modulates NSC activation/quiescence dynamics in the adult SEZ. In conjunction, these results are consistent with the P2Y_13_ receptor acting as a gatekeeper between NSC quiescence and activation, of this cell population in the adult SEZ *in vivo*.

**Figure 4.**
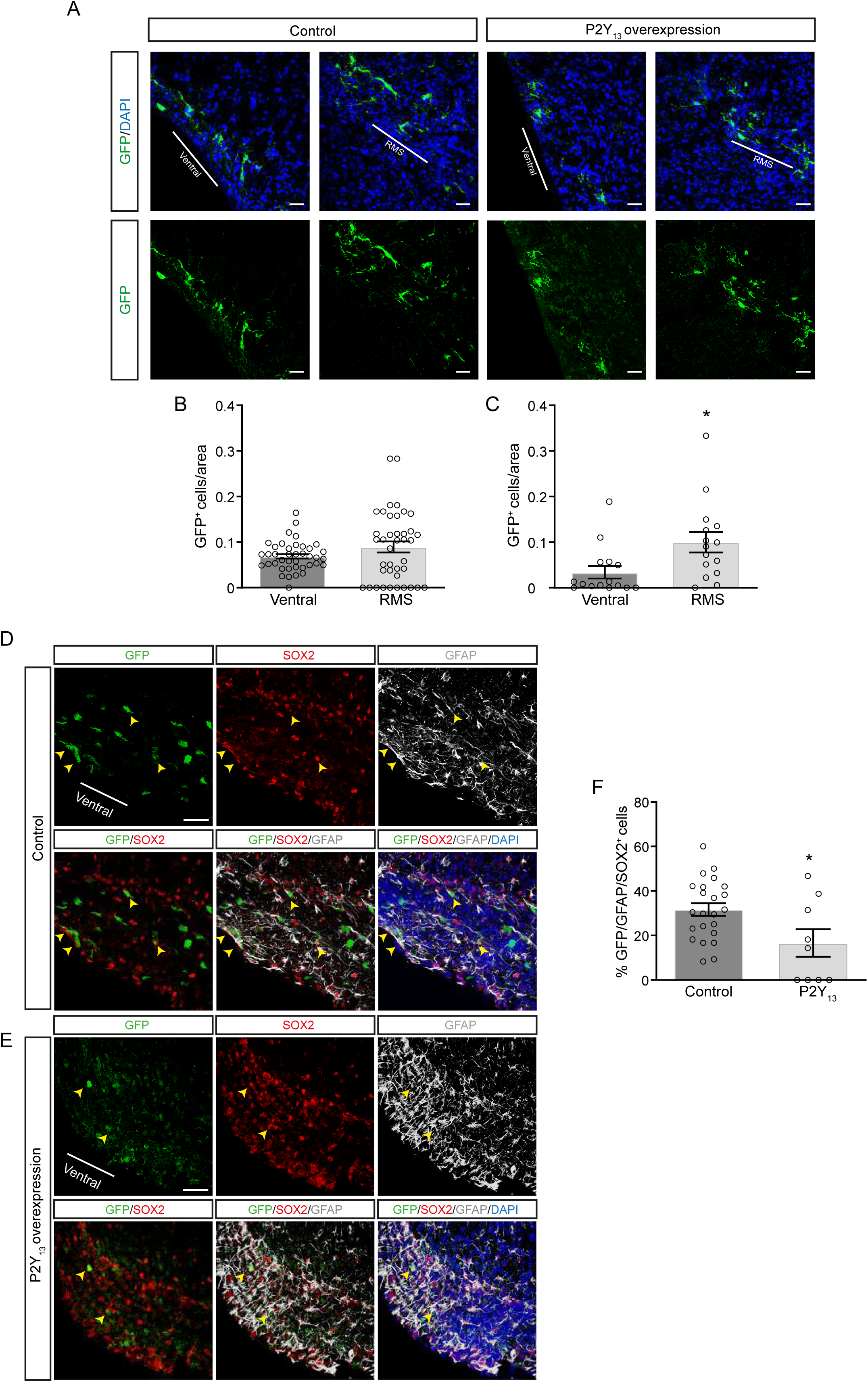
Local overexpression of P2Y_13_ receptors augments the proportion of cells in the RMS and decreases the number of NSCs remaining in the ventral wall of the SEZ. **A.** Effect of P2Y_13_ on SEZ dynamics, with local overexpression promoting more cells entering the RMS and less cells remaining in the SEZ (Scale bar 50 µm). **B** Quantification of GFP positive cells in the ventral wall of the SEZ and RMS following LV-GFP injection (n=4). **C** Quantification of GFP^+^ cells in the ventral wall of the SEZ and RMS following LV-P2Y_13_-GFP injections (n=4, scale bar 50 µm). **D-E.** Effect of local overexpression on the remaining GFAP^+^ (white)/SOX2^+^(red)/GFP^+^ (Green) cells in the SEZ after LV-GFP or LV-P2Y_13_-GFP injections (n=4, scale bar 50 µM). **F.** Quantification of GFAP^+^/SOX2^+^/GFP^+^ cells in the ventral wall of the SEZ following LV-GFP or LV-P2Y_13_-GFP injections (n=4). All graphs show the mean ±SEM: *p<0.05 (T-test).

### P2Y_13_ activity induces gene expression changes associated to activation and lineage progression *in vivo*

To gain insight into as to how P2Y_13_ activity may modulate adult NSC behavior, we sorted GFP^+^ cells within the neurogenic lineage progression from the SEZ 7 dpi of either LV-GFP (control samples) or LV-P2Y_13_-IRES-GFP (P2Y_13_ samples), which were then analyzed in RNA-seq experiments (**Figure 5A and S5**). A principal component analysis (PCA) revealed that both control and P2Y_13_ samples clustered together within their group (**Figure 5B**), and that cells transduced with LV-P2Y_13_-GFP had 1,432 upregulated and 59 downregulated genes relative to the control samples (**Figure 5E**). The transcriptome profile exhibited by the P2Y_13_ overexpressing cells revealed intriguing potential mechanisms by which the P2Y_13_ receptor may influence NSCs activation, mode of division and differentiation. Specifically, genes associated with activation and lineage progression were found to be upregulated, while conversely, markers of quiescence and, interestingly, self-renewal, were found to be downregulated. (**Figure 5D, E**). Specifically, we highlight the P2Y_13_-dependent upregulation of genes like the Colony stimulating factor 1 receptor (*Csf1r*) that is known to negatively regulate self-renewal and to promote neuronal differentiation in embryonic cortical NSCs (Nandi *et al*, 2012). Likewise, the upregulation of growth factor receptors was also observed and of some proteins directly related to NSC activation, like insulin growth factor 1 (*Igf1)*, the delta catalytic subunit of the PI3 Kinase (*Pik3cd*) and the Mitogen-activated protein kinase 14 (*Mapk14*) (de Morree & Rando, 2023). Other interesting genes upregulated by P2Y_13_ overexpression were galectin 3 (*Lgals3*) and its related gene galectin 3-binding protein (*Lgals3bp*), recently proposed to enhance astrocyte plasticity by leading to their activation and proliferation, and to neurosphere formation in several cortical-associated pathologies (Sirko *et al*, 2023). Another gene involved in neuronal activation and differentiation that was positively regulated by P2Y_13_ activity was *Stat6* (Bhattarai *et al*, 2016). Finally, an increase in the activity of the proteasome would be associated with the upregulation of *Psmb8, 9* and *10*, the preferred proteolytic pathway in activated NSCs, whereas lysosomal degradation is favored in qNSCs (Blasco-Chamarro & Farinas, 2023; Urban *et al*, 2019).

**Figure 5.**
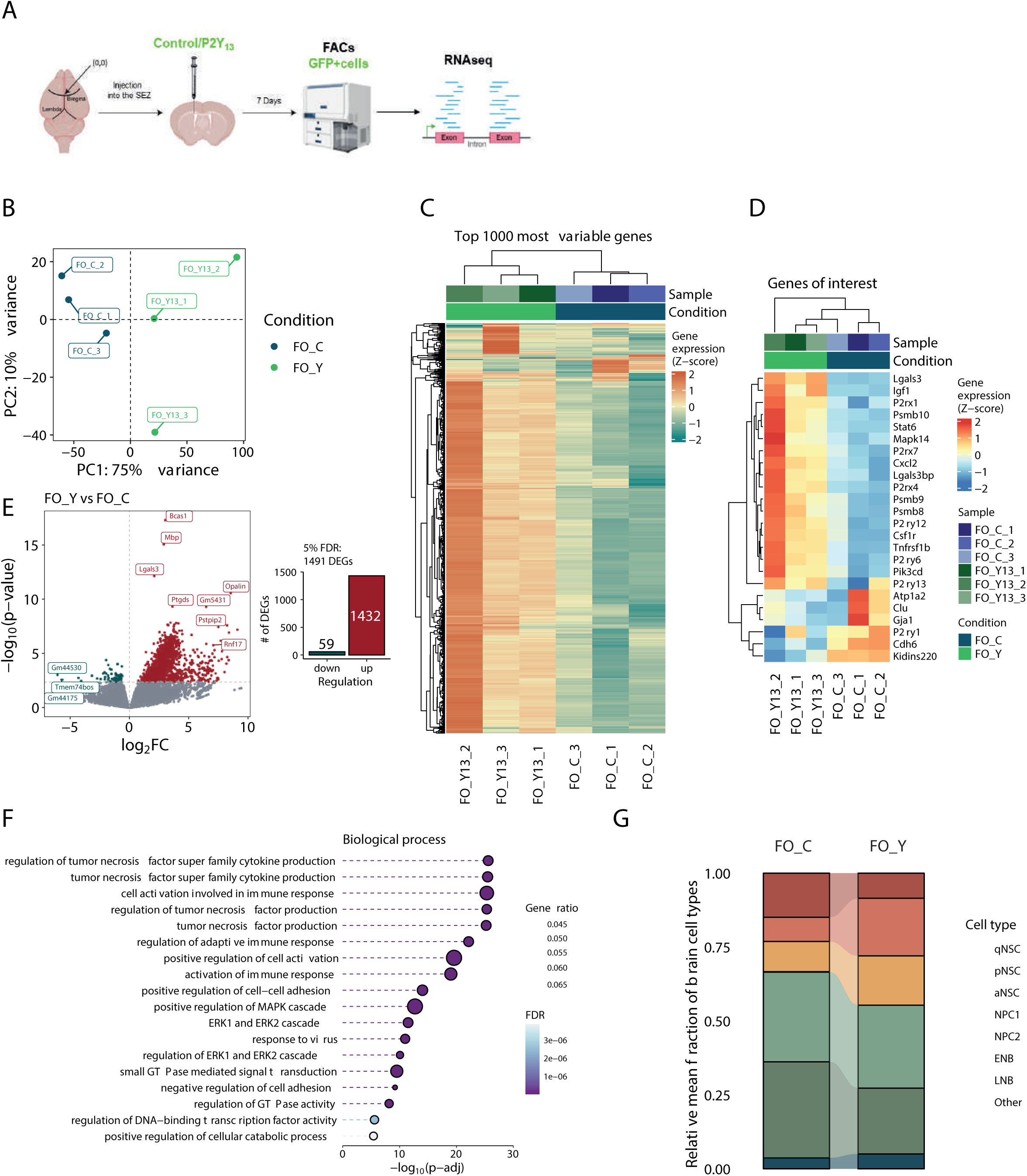
Transcriptomic analysis of the RNA-seq data obtained from P2Y_13_ overexpressing cells *in vivo*. **A.** Scheme of the experimental design: right panel created with BioRender.com. **B.** A principal component analysis (PCA) of the most variably expressed (top 3,000) genes of the samples profiled, colored by condition: control (*FO_C*, blue), P2Y_13_ overexpression (*FO_Y*, green). **C.** Heat-map of the top 1,000 most variably expressed genes across the samples profiled. The expression of each gene was scaled as a z-score, and the genes and sample labels were sorted by hierarchical clustering. **D.** Heat-map of the genes associated with the activation/quiescence equilibrium and self-renewal. The expression of each gene was scaled as a z-score, and the genes and sample labels were sorted by hierarchical clustering. **E.** Volcano plot showing the differential expression of genes between P2Y_13_ overexpressing and control samples (n=3, PY_13_ and 3 controls). Differentially expressed genes (DEGs, adjusted p-value <0.05) are in red and green when upregulated or downregulated in the P2Y_13_ overexpressing samples, respectively. The bar plot represents the number of DEGs. **F.** Overrepresentation of gene ontology (GO) terms for biological processes from the DEGs (adjusted p-value <0.05) between the P2Y_13_ overexpressing and control samples. A customized selection (18) of significant GO terms with the highest gene ratio (top 100) is displayed for clearer representation. The terms are ordered by significance, representing their adjusted p-value (x-axis), and the expressed genes were used as the background in this analysis. **G.** Stacked bar plot of the inferred cellular composition of neural progenitors for each condition. The cell deconvolutional analysis was carried out using CIBERSORTx, and the gene signatures of the neural progenitors were retrieved from the data generated in Belenguer et al., 2021.

Several genes associated with quiescence or self-renewal were negatively regulated by overexpression of the P2Y_13_ receptor, such as *Kidins 220* that has been associated with the promotion of gliogenic self-renewal divisions to maintain neurosphere formation (Del Puerto *et al*, 2023). We also observed the downregulation of several cadherins associated with qNSC populations (Morizur *et al*, 2018), such as *Cdh6.* Finally, several markers typically associated with qNSCs were also downregulated, like *Clu*, *Gja1* or *Atp1a2* (Dulken *et al*., 2017). An interesting issue was the regulation of the purinergic system itself, as P2Y_13_ overexpression also upregulated several other receptors, some of them positively associated with neuronal differentiation, survival and synapse formation like P2X4 and P2X7 (del Puerto *et al*., 2012; Diaz-Hernandez *et al*, 2008). Strikingly, the only purinergic receptor downregulated was the P2Y_1_ receptor, known to be involved in the maintenance of stemness in embryonic radial glia (Weissman *et al*., 2004).

When gene ontology (GO) terms were analyzed (**Figure 5F**), some genes upregulated by the local overexpression of P2Y_13_ receptors were associated with different signaling pathways implicated in the modulation of quiescence and activation, as well as in neuronal differentiation. Amongst these signaling pathways was the modulation of GTPase signal transduction and activity, which is known to directly modulate NSC quiescence (Chavali *et al*, 2018). Likewise, P2Y_13_ overexpression modulated several terms positively associated with activation and transcription, while negatively associated with cell-adhesion, which favors NSC activation to the detriment of quiescence (Blasco-Chamarro & Farinas, 2023; Morizur *et al*., 2018; Urban *et al*., 2019). In addition, terms related to the modulation of the Mitogen-activated protein kinase (MAPK) cascade and within it, to the extracellular signal-regulated kinase (ERK) pathway, were also recognized, which are known to be involved in the activation of a proneural genetic switch in embryonic NSCs (Li *et al*, 2014). This was in line with the P2Y_13_-dependent activation of ERK proteins previously shown to be involved in neuronal survival (Ortega *et al*., 2011b; Ortega *et al*, 2008). Furthermore, the abundance of terms associated with the modulation of the immune response was notable, involved in the balance between activation and the return to quiescence of NSCs (Belenguer *et al*., 2021b; Kyritsis *et al*, 2012).

Finally, the transcriptomic profile of the isolated GFP^+^ cells was assessed, bearing in mind the molecular signatures of the different neural populations within the NSC-derived lineage progression described previously (Belenguer *et al*., 2021b). Our deconvolution analysis showed that although the qNSC profile was not detected in our comparisons, P2Y_13_ overexpression produced a more active phenotype, with a higher percentage of our transcriptomes associated with aNSC and NPCs (**Figure 5G**). Interestingly, we detected similar fractions corresponding to early NBs but fewer to late NBs, which is consistent with the stronger activation observed *in vivo* and the probable presence of these cells in the RMS when the GFP^+^ cells were isolated from the ventral wall of the SEZ in our experiments. These findings not only confirm our in vivo results but also suggest broader implications for understanding the balance between stem cell maintenance and differentiation in the adult brain.

### P2Y_13_ activity modulates the balance between NSC quiescence and activation

Our data suggested the possibility that P2Y_13_ receptor activity is associated with an activation of NSCs linked to their subsequent lineage progression towards symmetrical neuronal differentiation. Therefore, we next sought to confirm the specific effect of the P2Y_13_ receptor on NSC activation/quiescence and self-renewal at a single cell level by exploiting a method developed in house that enables single cell tracking of NSCs, isolated from their neurogenic niche signals and in the absence of added mitogens, and their individual progeny in real-time, defining their lineage progression trees. These lineage trees capture the behavior of NSCs and allow the tracking of NSC fate decisions at the single cell level to assess the balance between asymmetric and symmetric divisions, self-renewal and differentiation, cell death or survival, and quiescence and onset of activation. Importantly, using these methods, we have reportedly demonstrated that complex trees (having undergone 4-5 rounds of division) are generated precisely by the activation of *bona fide* NSCs, whereas less complex trees (<3 rounds of division) are produced mainly by TAPs and early NBs, with trees reflecting ≥6 rounds of division being particularly scarce in isolated primary cultures (Costa et al, 2011; Ortega et al., 2011a; Ponti et al., 2013)(Costa et al., 2011; Gomez-Villafuertes et al., 2017; Ortega et al., 2013a; Ortega and Costa, 2016; Ortega et al., 2011a). Moreover, since isolated NSCs maintain their neurogenic potential despite the absence of signals from their niche, this approach also represents a unique tool to study the effects of specific signaling molecules and to search for markers that could distinguish the heterogeneous states of NSCs (Bicker et al., 2017; Domingo-Muelas et al., 2023; Farrukh et al., 2017; Garcia-Gonzalez et al., 2016; Ortega et al., 2013b). Thus, primary cultures of the adult ventral SEZ, either under control conditions or exposed to the P2Y_13_ receptor agonist 2MeSADP, were monitored by time-lapse video-microscopy for single-cell tracking followed by post-imaging ICC. Indeed, the proportion of clones that achieved greater complexity through divisions (4 and 5 rounds of amplifying divisions) after 5 DIV significantly increased upon activation of the P2Y_13_ receptor (7.88 ± 2.23% and 5.43 ± 1.77%, respectively) relative to the control conditions (4.50 ± 1.25% and 0.81 ± 0.50%, respectively; **Figure 6A-D**). This increase could reflect more NSCs being activated and undergoing lineage progression, in line with our *in vivo* experiments, enhanced proliferation within the neurogenic trees, or enhanced survival of neural progenitors, thereby provoking the establishment of more complex trees. However, single cell tracking revealed that the survival rate within the neurogenic trees was not altered (**Figure 6E; Figure S6**). Moreover, the increase in the number of complex trees was not a consequence of a decrease in the number of clones that underwent 1-3 rounds of division, suggesting that the proliferative capacity within those clones was not enhanced, but rather there was an increase in the number of progenitors being activated and progressing towards neurogenesis (**Figure 6C; Figure S6**). Finally, to further confirm a specific effect of the P2Y_13_ receptor on NSC activation as opposed to proliferation rate, primary cultures of the adult SEZ were exposed to the specific P2Y_13_ agonist, the 2MeSADP and analyzed 6 days after plating. There was only a minimal increase in the number of βIII-tubulin^+^ neurons detected (84.50 ± 1.84% in 2MeSADP and 77.30 ± 3.26% in control condition; **Figure S7A**), consistent with an increase in NSC activation rather than a global effect on proliferation, especially considering that the GFAP^+^/SOX2^+^ NSC population represents only a 16.51 ± 1.77% and 5.95 ±1.34% of the whole cell population on the first and second day *in vitro*, respectively (**Figure S7B**). Thus, our data suggest that the activation of the P2Y_13_ receptor triggered by 2MeSADP instructs NSCs to set out their program of lineage progression.

**Figure 6.**
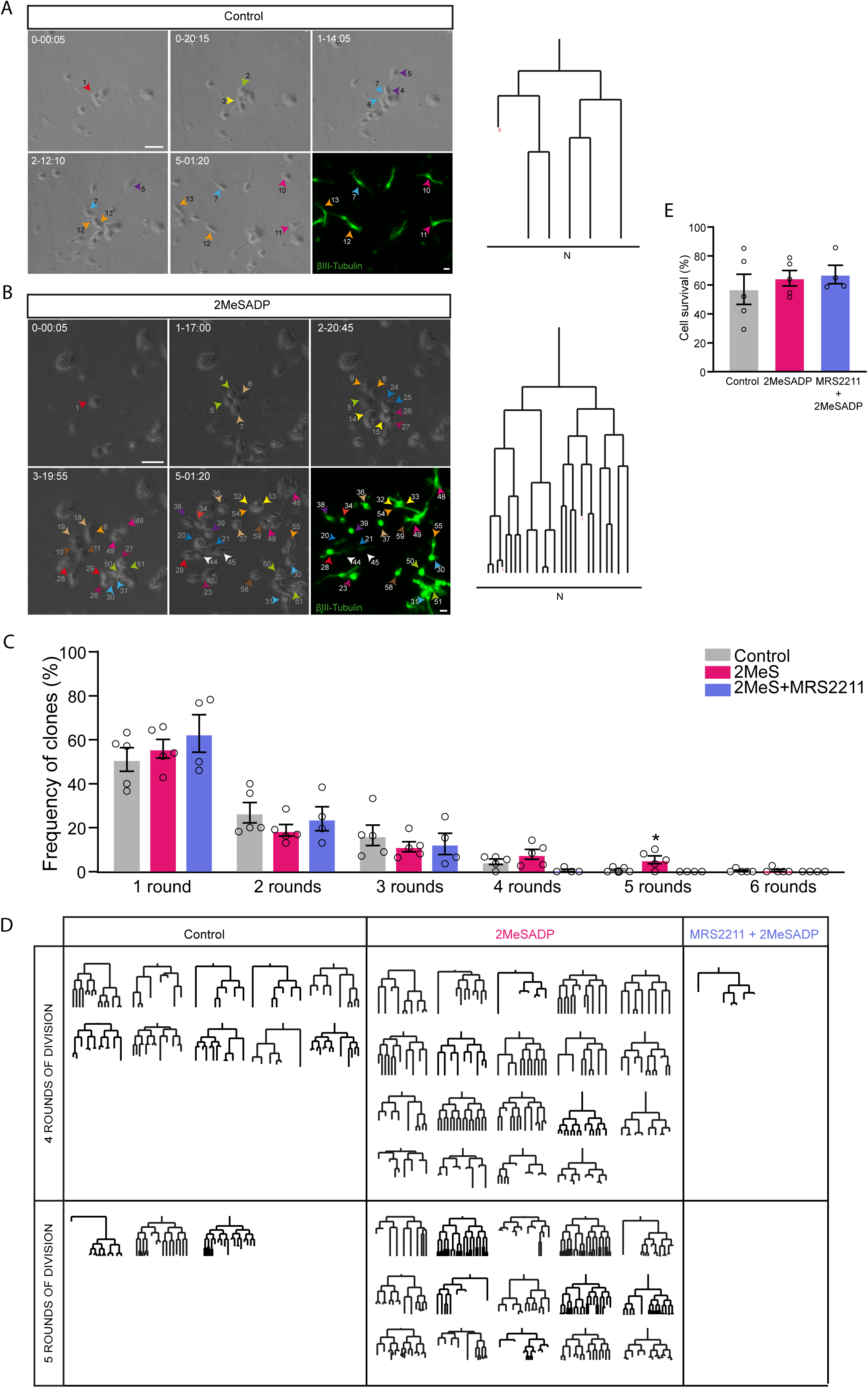
Neurogenic trees from the ventral wall of the SEZ tracked in culture following treatment with the P2Y_13_ agonist 2MesADP, in the presence or absence of the MRS2211 antagonist. **A** Representative symmetric neurogenic tree obtained in control conditions after 6 days in culture (N, neuron; X, cell death). **B** Complex symmetric neurogenic trees of 5 rounds of division obtained in the presence of 2MeSADP after 6 days in culture (N, neuron; X, cell death). The phase contrast images in both **A** and **B** depict the lineage progression in the live imaging experiment (day-hour-min). The last images show post-imaging immunocytochemistry of the neuroblast progeny (βIII-tubulin in green) Scale bar 30 µm. **C** Clones undergoing 1-6 rounds of division in the live imaging experiments (n=5) (n=4 for MRS2211). **D** Summary of all the clones tracked undergoing 4 or 5 rounds of division in our live imaging experiments, either in control conditions, or when exposed to 2MeSADP or 2MeSADP + MRS2211 (n=5) (n=4 for MRS2211). **E** Cell survival in the lineage trees (n=5). In all cases the progeny generated were identified by post-imaging immunocytochemistry. All graphs show the mean ±SEM: *p<0.05, and **p<0.01 (ANOVA with a Tukey’s post-test).

When cultures were maintained in the presence of both 2MeSADP and MRS2211 (**Figure 6C-E**) the antagonist reverted the effect of 2MeSADP on NSC activation and the generation of complex trees (4 and 5 rounds of division), which were barely detected when MRS2211 was added to the cultures (**Figure 6C,D**) while leaving the survival rate within the clones unaffected. It is important to note that the reduction in the number of complex trees (<3 rounds of division) was not due to reduced proliferation but rather, there were more cells that remained quiescent throughout the experiment (see below). Together these experiments point to a potential role of the P2Y_13_ receptor as an active regulator of the balance between NSC quiescence and activation in the adult SEZ.

As previously mentioned, adult neurogenic niches are known to be exposed to the tonic release of several neurotransmitters (Berg *et al*, 2013), including ATP and other related nucleotides (Lin *et al*., 2007; Young *et al*., 2011). To assess if the tonic activation of P2Y_13_ might influence its effects on NSC progression, time-lapse video-microscopy experiments were performed in the presence of the P2Y_13_ antagonist MRS2211 to counteract the potential tonic activation of this receptor in our control experiments. Consistent with the tonic activation of the P2Y_13_ in the adult SEZ, exposing primary SEZ cultures to MRS2211 significantly reduced the number of clones that went beyond 3 rounds of division when tracked by live imaging (**Figure S7C**).

As suggested by our *in vivo* experiments, we then addressed whether the P2Y_13_ receptor might also affect other inherent hallmarks of adult NSCs, i.e. their ability to generate new copies of themselves through asymmetric division, also known as self-renewal events (Obernier & Alvarez-Buylla, 2019). Live time-lapse video-microscopy followed by post-imaging ICC enables NSC progeny to be identified, classifying the lineage trees as either symmetric neurogenic trees (i.e. generating only NBs) or asymmetric trees comprised of NBs and GFAP^+^ astrocytes and/or GFAP^+^/SOX2^+^ NSCs, and involving a self-renewal event (Costa *et al*., 2011; Ortega *et al*, 2013a; Ortega *et al*, 2011a). Exposure of the cells in culture to 2MeSADP significantly reduced the proportion of asymmetric lineage trees tracked (10.6 ± 3.7% in 2MeSADP and 23.4 ± 12.84% in control condition; **Figure 7A**), indicating a dramatic reduction in the number of self-renewal events upon P2Y_13_ activation. Strikingly, not only did the P2Y_13_ antagonist MRS2211 fail to impede the effect of 2MeSADP but it further reduced the proportion of asymmetric trees to 4.54 ± 4.54% (**Figure 7A**). Considering that MRS2211 does not affect cell survival within the lineage trees, this effect could potentially point to a non-specific effect of the P2Y_13_ receptor on the regulation of self-renewal events. To confirm this possibility, we monitored the NSCs for three days to evaluate the dynamics of the early lineage trees prior to neuronal differentiation. To verify that these trees were in fact NSCs-derived trees, enhancing the specificity of the results, only clone-founder NSCs that gave rise to asymmetric lineage trees containing new copies of GFAP^+^/SOX2^+^ progeny were considered. This is relevant as after only three days of imaging, it is not clear if symmetric trees formed entirely by GFAP^-^/SOX2^-^ cells are founded by a NSC or by a TAP, and it is not possible to discern at these early time points whether these trees will progress towards higher complexity (≥4 rounds of division), potentially a distinguishing hallmark. Post-imaging ICC allowed us to classify the initial NSCs (24 h after plating) into three experimental subgroups: the “Initial” group of all NSCs present at the beginning of the live imaging experiment, the “Quiescent” group of initial NSCs that did not advance through lineage progression and remained quiescent throughout the experiment, and the “Clones” group of initial NSCs that embarked on lineage progression and generated asymmetric clonal trees (**Figure 7B**). Importantly, live imaging experiments clarified the apparent contradictory effect between 2MeSADP and MRS2211. While 2MeSADP induced fast activation and differentiation, decreasing the number of asymmetric NSC trees, MRS2211 did not reduce the asymmetry in the same way but rather, it left NSCs quiescent throughout the experiments (**Figure 7B**). This was particularly evident as the proportion of initial (0.39 ± 0.13%) and quiescent NSCs (0.31 ± 0.04%) in the culture almost overlapped in the presence of MRS2211, with only a minimal fraction of NSCs entering lineage progression. To further disentangle the mechanisms of this effect, it is worth briefly considering the standard behavior of adult SEZ-derived NSCs when isolated *in vitro* and maintained in the absence of mitogenic factors (Costa *et al*., 2011; Ortega *et al*., 2013a; Ortega *et al*., 2011a) (**Figure 7C**). During the first 1-2 days *in vitro*, the NSC cell cycle is slow prior to division, subsequently undergoing one or two divisions. These are initially symmetric astrogliogenic divisions, which lead to the generation of “fast dividing-astroglia” (i.e. GFAP^+^/SOX2^+^ cells) that go on to generate TAPs and the rest of the lineage. As a result, there is an initial expansion of the NSC pool in the first 24-48 h. Moreover, it is common that one of the branches of this NSC-based lineage tree gives rise to quiescent GFAP^+^/SOX2^+^ cells as a self-renewal branch (see model in **Figure 7C**, “Control”). This characteristic behavior was altered in the presence of 2MeSADP, which accelerated the NSC cell cycle, particularly the first rounds of division (i.e. where fast-dividing astroglia are generated; **Figure 7D, E**). This stimulation produces differences in the “Initial” group (**Figure 7B**) as exposure to 2MeSADP drives NSCs transition to TAPs within the first 24 h of live imaging. When combined with fewer self-renewal events, P2Y_13_ receptor activation reduced the overall number of asymmetric lineage trees (**Figure 7A, C-E,** “2MeSADP”). Conversely, MRS2211 instructed NSCs to remain mostly quiescent throughout, leading to a drastic reduction of NSCs in the “Initial” group as they did not undergo expansion during the first 24-48 h, as described previously (**Figure 7B, C** “MRS2211+2MeSADP”). Accordingly, only very few of the NSCs embark on lineage progression and confusingly, there is an apparent reduction in self-renewal events. Taken together, these experiments further confirm our *in vivo* results and postulate that the purinergic signaling, acting through the P2Y_13_ receptor, constitutes not only a key regulator in the balance between activation and quiescence of SEZ NSCs, but, remarkably, also links its activity to the mode of division of NSCs once the lineage progression is initiated, reducing the number of their self-renewal events in favor of differentiative divisions.

**Figure 7.**
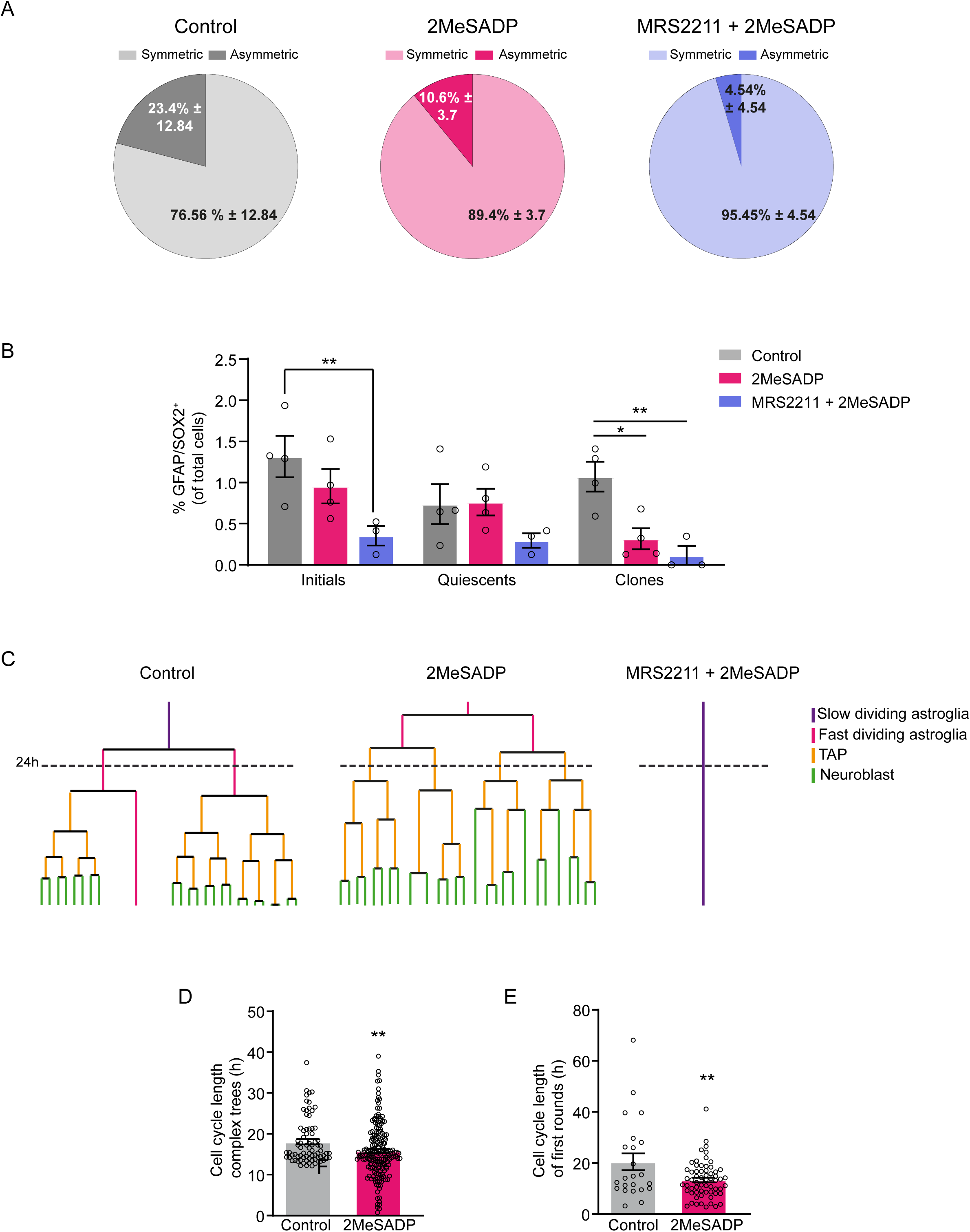
Effect of the P2Y_13_ receptor on NSC self-renewal. **A**. Quantification of the symmetric lineage trees (generating only neuroblasts) and the asymmetric lineage trees (generating neuroblasts and new NSCs through self-renewal events) in control conditions, or on exposure to 2MeSADP or 2MeSADP + MRS2211 (n=4). **B.** Proportion of GFAP^+^/SOX2^+^ NSCs relative to the total number of cells in the culture according to the following labels: “Initial” indicates the proportion of GFAP^+^/SOX2^+^ in the culture and the “Quiescent” cells are the fraction of the initial cells that remain quiescent throughout the live imaging experiments. The “Clones” reflect the fraction of the initial cells that undergo lineage progression (Control and 2MeSADP n=5 and 2MeSADP+MRS2211 n = 4). **C.** The models represent the stereotypic behavior of NSCs under control conditions and when exposed to 2MeSADP or 2MeSADP+MRS211. Note how 2MeSADP increases the speed in the cell cycle while impeding self-renewal divisions. By contrast, MRS2211 instructs NSCs to remain quiescent. **D.** Cell Cycle length within the complex trees tracked (≥4 rounds of division). **E**. Cell Cycle length for the first division within the complex trees tracked (≥4 rounds of division, n= 5). Note that this first division is normally associated with slow-dividing astroglia, yet 2MeSADP significantly increase the speed of the cell cycle (n=5). All the graphs show the mean ±SEM: *p<0.05, **p<0.01 (T-test for **D, E** and ANOVA with a Tukeys post-test for **B**).

## DISCUSSION

In this study, evidence is presented that the purinergic P2Y_13_ receptor conspicuously labels NSCs in the adult SEZ and more specifically, that it can be used to differentiate between qNSCs and aNSCs. Although the presence of P2Y receptors in the adult SEZ has been explored, previous studies employed either classic neurosphere cultures or addressed the entire cell landscape within the intact SEZ *in vivo*. In addition, the lack of specific agonists and antagonists have made it difficult to gain accurate information regarding the specific effects of P2Y receptor expression as the neural lineage progresses (Ali *et al*, 2021; Grimm *et al*, 2009; Mishra *et al*, 2006). However, the data obtained here suggests a preferential presence of the P2Y_13_ receptor in the ventral wall of the SEZ, an area populated by NSCs more prone to induce neurogenesis (Colak *et al*., 2008; Ortega *et al*., 2013b). Moreover, the P2Y_13_ receptor specifically localizes to the qNSC pool, situating the P2Y_13_ receptor as a potential marker to fate-map the contribution of the qNSC to lineage progression. Its restricted expression opens the possibility of using the P2Y_13_ receptor as a valid marker, overcoming the lack of means to effectively discern between mature astrocytes and qNSCs in humans (Bhaduri *et al*, 2020). This is an interesting issue that will require confirmation in human tissue. Furthermore, the data serves to reinforce the versatility of the existing methods for the isolation of this population, based on an intricate combination of markers or in highly resource-consuming single-cell analysis.(Basak *et al*., 2018; Belenguer *et al*., 2021a; Codega *et al*., 2014; Dulken *et al*., 2017; Fischer *et al*, 2011; Giachino *et al*., 2014b; Llorens-Bobadilla *et al*., 2015; Mich *et al*, 2014; Morizur *et al*., 2018; Pastrana *et al*, 2009).

Importantly, the P2Y_13_ receptor appears to regulate the balance between activation and quiescence in the NSC pool. In addition, the P2Y_13_ receptor seems to control one of the main hallmarks of somatic stem cells, their ability to generate new copies of themselves through self-renewal (Lim & Alvarez-Buylla, 2014; Obernier & Alvarez-Buylla, 2019).

Thus, the modulation of neurogenesis in the DG by P2Y_13_ expressed in microglia reported previously (Stefani *et al*., 2018) is now complemented by the influence of the P2Y_13_ receptor on the neural lineage in the SEZ. Moreover, our data highlight the role of purinergic metabotropic neurotransmission as part of the modulatory program of G-protein coupled receptors in adult neurogenesis (Doze & Perez, 2012; Lin *et al*, 2024). Likewise, our results add to the growing evidence as to how neurotransmitter activity regulates NSC lineage progression (Berg *et al*., 2013; Giachino *et al*, 2014a; Song *et al*, 2012; Tong *et al*, 2014; Trinchero *et al*, 2021). Furthermore, the increase in the intracellular calcium levels observed upon activation of the P2Y_13_ receptor is consistent with the calcium-dependent modulation of the transition between quiescence and activation in the SEZ (Gengatharan *et al*, 2021).

Extracellular ATP-mediated variations in cellular calcium have also been said to be involved in regulating the proliferation and activation of embryonic stem cells and human mesenchymal stem cells (Heo & Han, 2006; Kawano *et al*, 2006). However, as indicated in terms of P2Y receptor expression in the SEZ, the studies performed to date address the impact of purinergic neurotransmission on adult neurogenesis using a broad approach, with little specificity in terms of the agonists and antagonists employed. Moreover, most of these studies rely on either population analyses performed *in vivo* or on the study of isolated cells cultured in the presence of mitogens as neurospheres (Grimm *et al*., 2009; Mishra *et al*., 2006). Thus, in addition to the confounding effects exerted by mitogens on NSC behavior is added the loss of information that occurs in experiments based on end-point analysis (Costa *et al*., 2011; Ortega & Costa, 2016; Pastrana *et al*, 2011; Schroeder, 2008). For instance, the activation of a relatively small population of qNSCs in the SEZ or of the vast number of cells of the whole lineage isolated *in vitro*, can be mistakenly detected as a transient increase in cell proliferation. Conversely, the continuous live imaging and single cell tracking of NSCs isolated from the adult SEZ allows heterogeneous cell behaviors, cell fate decisions or cell death to be studied within a clone (Ortega & Costa, 2016). As such, the specific effect of the P2Y_13_ receptors on both activation and self-renewal can be disentangled, showing how its activity enhances NSC activation and generates new neurons symmetrically, while its inhibition retains NSCs in a quiescent state.

Moreover, experiments based solely on population analyses that are performed *in vivo* make it unfeasible to rule out the simultaneous presence of different elements of the purinergic system in the adult SEZ, these potentially acting in concert to fine-tune different aspects of neurogenic progression. Indeed, we already described this phenomenon in other neurogenic niches (Paniagua-Herranz *et al*., 2020b) By contrast, live imaging performed on cells isolated from the niche allowed us to address how pharmacological modulation of the P2Y_13_ receptor affects these issues, at least *in vitro*.

The data presented also points to the potential involvement of MAPK/ERK-associated signaling pathways, as well as the modulation of the immunogenic/inflammatory response, as molecular mechanisms triggered in response to P2Y_13_ activation. The participation of ERK signaling is consistent with the known activation of proneural genetic programs in Radial Glial NSCs, modulating Ascl1 activity and promoting neuronal differentiation (Li *et al*., 2014). Moreover, we previously described how the P2Y_13_ receptor modulates ERK signaling to promote cerebellar granule neuron survival when faced with glutamate-induced excitotoxicity, genotoxic stress, oxidative stress and trophic factor withdrawal (Espada *et al*., 2010; Morente *et al*., 2014; Ortega *et al*., 2011b; Ortega *et al*., 2008). Importantly, the P2Y_13_ receptor affords this neuroprotection by interacting with intracellular signaling pathways modulated by trophic factors like the Insulin Growth Factor (IGF) receptor/PI3K axis, also known to participate in adult neurogenesis (Yuan *et al*, 2015) and seen to been positively regulated in the transcriptomic analysis performed here. Moreover, the modulation of the immunogenic response observed upon P2Y_13_ activation correlates with the recent observation on the role of these molecules in controlling the balance between activation and quiescence (Belenguer *et al*., 2021b; Blasco-Chamarro & Farinas, 2023), influencing both the transition from a quiescent to an alert state, as well as reversion to a dormant state. Thus, our results contribute to the interesting debate on how the proinflammatory/immunogenic response unfolds in the absence of any existing injury or inflammation, and how it may regulate NSC lineage progression within the adult neurogenic niches (Belenguer *et al*., 2021b; Kalamakis *et al*, 2019; Kyritsis *et al*., 2012).

Finally, by regulating the balance between quiescence/activation, the P2Y_13_ receptor may represent a starting point for the design of strategies to mitigate pathological conditions. For instance, insufficient quiescence can promote tumor formation or depletion of the NSC pool (Urban & Cheung, 2021). Conversely, excessive quiescence can lead to a loss of tissue homeostasis due to more limited generation of new cells (Cheung & Rando, 2013). This is similar to the situation in aging, where a significant fraction of NSCs remains in a quiescent state, precisely due to the expression of various pro-inflammatory transcription factors that enhance quiescence in the niche where they exist. These cells also exhibit a loss of regenerative capacity and greater resistance to activation, even after injury (Baruch *et al*, 2014; Kalamakis *et al*., 2019; Villeda *et al*, 2011). Therefore, the P2Y_13_ receptor emerges as a very promising tool to force the activation of NSCs in cases of neuronal death related to aging.

## METHODS

### Ethics Statement

All animal procedures were carried out at the UCM in accordance with European (2010/63/EU) and Spanish (RD1201/ 2005; RD 53/2013) regulations and following the guidelines of the International Council for the Laboratory Animal Science. The experimental protocols were approved by both the Committee for Animal Experimentation of the UCM and of the Regional Government of Madrid.

### Primary cell cultures and drugs

Primary cultures of NSCs from the mouse SEZ were established following the procedure described previously (Costa *et al*., 2011; Ortega *et al*., 2013a; Ortega *et al*., 2011a). Briefly, the lateral wall of the lateral ventricle of young adult (8-12 weeks) C57/B16 mice was enzymatically dissociated in 0.68 mg/mL trypsin and 0.7 mg/mL hyaluronic acid (both from Sigma-Aldrich) prepared in Hanks’ Balanced Salt Solution (HBSS: Gibco), with 2 mM glucose and buffered with 15 mM HEPES at 37 °C for 30 minutes. Enzymatic dissociation was stopped by adding an equal volume of a cold solution of 4% bovine serum albumin (BSA: Sigma-Aldrich) in Earle’s Balanced Salt Solution (EBSS: Gibco) buffered with 20 mM HEPES (Gibco). The cells were then centrifuged at 300 *g* for 5 minutes at 4 °C, and the cell pellet was re-suspended in an ice-cold solution of 0.9 M sucrose (Sigma-Aldrich) in 0.5x HBSS and centrifuged at 700 *g* for 10 minutes at 4 °C. The sediment was then re-suspended in 1 mL of a cold 4% BSA solution in EBSS buffered with 20 mM HEPES and then another 12 mL of the same solution was added. After centrifuging at 4 °C for 7 min at 400 *g* and the sediment obtained was finally re-suspended in DMEM-F12 culture medium supplemented with 2% B27, 100 U/mL penicillin, 100 μg/mL streptomycin and 0.5 mM glutamine (all from Gibco). The resulting cell suspension was seeded in a P24 multiwell plate coated with poly-D-Lysine (PDL) (Sigma) in sterile ultrapure water at 0.1 mg/mL and maintained at 37 °C in a humidified atmosphere containing 8% CO_2_. The 2MeSADP, MRS221 and MRS2179 were purchased from Tocris Bioscience.

### Time-lapse videomicroscopy

Live imaging was performed using a NIKON TE-2000 microscope as described elsewhere (Gomez-Villafuertes *et al*, 2017; Paniagua-Herranz *et al*, 2020a) at a constant temperature of 37 °C and in 8% CO_2_. Images were obtained every 5 minutes over 4-5 days using a long-distance 20x phase contrast objective (Nikon), a ZYLA camera (ANDOR) and the 4.7/NIS-elements software from NIKON. The analysis of the images was performed with tTt software (The Tracking Tool: (Hilsenbeck *et al*, 2016), through which the cells of interest were tracked at the selected position and lineage trees were reconstructed. The identity of the progeny was determined by post-imaging immunocytochemistry (ICC) and movies were assembled using ImageJ 1.42q (National Institute of Health, USA).

### RT-PCR and quantitative real-time PCR

Total RNA was obtained from SEZ tissue using the SpeedToold Total RNA Extraction kit (Biotools), according to the manufacturer’s instructions. After quantification, 1 μg of the purified RNA was reversed transcribed and then quantitative real-time PCR (qPCR) was performed with the LuminoCt qPCR Ready Mix Amplification Kit (Sigma), using TaqMan MGB probes from (Roche Diagnostics, Barcelona, Spain). The qPCR assays were carried out using a StepOnePlus Real-Time PCR System (Applied Biosystems). The reaction proceeded in a polymerase activation phase for 20 seconds at 95 °C, followed by 40 cycles of amplification (denaturation phase for 1 second at 95 °C, and hybridization and elongation phase for 20 seconds at 60 °C). The expression of each transcript was quantified by the 2 double delta Ct (2^-ΔΔCt^) method, using GAPDH gene expression as an internal amplification control.

### RNA seq

#### • Ultra-low input RNA library preparation and sequencing

The SMART-Seq v4 (SSv4) Ultra Low Input RNA Kit for Sequencing (Takara #634888) was used to generate high-quality and full-length cDNA. Initially, RNA was purified from the total RNA using oligo(dT) attached to magnetic beads. The samples were amplified by LD-PCR over 18 cycles, followed by cDNA purification, controlling for average length and yield using Qubit 4 (Thermo Fisher) and Bioanalizer 2100 (Agilent), respectively. The amplified cDNA was used to pool the libraries prior to preparing them with the NEB Next® Ultra™ RNA Library Prep Kit, and they were then sequenced on an Illumina NovaSeq 6000 platform using paired-ends and with a read length of 150 bp, producing a total of 6 Gb of data (∼20 million read pairs per library).

#### • RNA-Seq data processing

The quality of the RNA-Seq data was controlled using FastQC software (v0.11.9) (S., 2010)before and after trimming, and the samples accepted were used for alignment. Paired-end reads were mapped to the mouse genome assembly (GRCm39) from the ENSEMBL database release 106 using STAR v2.7.9a (Dobin *et al*, 2013)retaining the uniquely mapped reads in correct pairs. Gene-level abundance was estimated using the gene counts option in the STAR software for the GENCODE vM29 annotation and using the primary assembly annotation gtf file. Downstream analyses was performed using the R v4.2.2 (Team, 2013)and Bioconductor v3.16 (Love *et al*, 2014)packages.

#### • RNA-Seq data analysis

##### - Quality control

Quality control of the sample similarity between conditions (P2Y13 versus Control) was based on a PCA and on hierarchical clustering of the transformed gene counts (variance stabilized). Previously, weakly expressed genes (i.e.: with a mean expression <1 read across all samples) were removed from further downstream analysis, retaining 27,681 genes. The PCA was performed on the top 3,000 variable genes using the DESeq2 package (v1.38.3: 5) (Love *et al*., 2014), a procedure that identified any batch effect prior to the differential expression analysis.

##### - Differential gene expression

Differential gene expression was analyzed on the P2Y13 samples using the DESeq2 package (v1.38.3: 5) (Love *et al*., 2014) applying the default parameters and using the *“∼ mutated*” model to contrast P2Y13 overexpression relative to the control samples. Genes with an adjusted p-value <0.05 were considered differentially expressed genes (DEGs).

##### - Gene ontology overrepresentation analysis

Overrepresentation of gene ontology (GO) terms for biological processes was performed with the DEGs using clusterProfiler v4.6.2 (Wu *et al*, 2021) and org.Mm.eg.db v3.16.0 (M, 2019), using expressed genes as the background.

##### - Cell deconvolution analyses

The CIBERTSORTx tool (Newman *et al*, 2019)was used to estimate the relative fraction of neural progenitor cells in each condition. As such, a limma-voom transformed matrix with the gene signatures of these specific cell types was obtained from data available elsewhere (Belenguer *et al*., 2021b)and used as a signature matrix. Following the recommendations in the documentation, the same normalization method was implemented on our raw gene expression data (exclusive to this analysis) to generate the mixture matrix. The limma v3.54.2 (Ritchie *et al*, 2015)voom (Law *et al*, 2014)ENSEMBL gene IDs were converted to gene symbols using GENCODE vM29 annotation and then, both matrices were anti-log transformed and submitted to the tool to impute the cell fractions, specifying: B-mode batch correction, 100 permutations and the absolute mode.

### Immunocytochemistry (ICC) and immunohistochemistry (IHC)

For ICC, cells were seeded on coverslips or on the bottom of the well, they were fixed with 4% paraformaldehyde (PFA) for 10 minutes and then blocked for 1h at room temperature (RmT) in ICC blocking solution: PBS containing 2% BSA and 0.2% Triton X-100 (v/v). For IHC, brain sections of the SEZ were obtained from mice that had been perfused transcardially with 4 % PFA. The brains were cryoprotected in a 30% sucrose solution in PBS for 48 hours at 4 °C and then embedded in O.C.T. (Sakura Finetek). Sagittal cryostat sections (20 μM: Leica CM1950) were then permeabilized and blocked for 1h at RmT in IHC blocking solution: PBS containing 0.2% Triton X-100 (v/v), 5% fetal bovine serum (FBS) and 1% BSA. Vibratome sections (70 μm thick) were obtained from brains embedded in 4% (w/v) agarose in PBS, permeabilized and blocked for 1 h floating at RmT in vibratome blocking solution: PBS containing 2% BSA and 0.5% Triton X-100 (v/v).

Both the cells and tissue sections were then incubated overnight at 4 °C with the primary antibodies: rabbit anti-P2Y_13_ (1:100, Alomone Labs Cat# APR-009), rabbit anti-P2Y_1_ (1:100, Alomone Labs Cat# APR-017), rabbit anti-SOX2 (1:100, ABclonal Cat# A0561), mouse anti-GFAP (1:200, Sigma-Aldrich Cat# G3893), mouse anti-βIII-Tubulin (1:800, Sigma-Aldrich Cat# T8660), guinea pig anti-(Doublecortin)DCX (1:400, Millipore Cat# AB2253), mouse anti-ASCL1 (1:100, BD Biosciences Cat# 556604), rabbit anti-KI67 (1:100, Fisher Scientific Cat# RM-9106-S), and chicken anti-GFP (1:400, AvesLab Cat# GFP-1020). After washing the sections three times with PBS/3% BSA (v/v) for 1h at RmT the cells/sections were probed with secondary antibodies: Alexa Fluor 488 goat anti-rabbit IgG (H+L: Cat# A-11008), Alexa Fluor 594 goat anti-rabbit IgG (H+L: Cat# A-11012), Alexa Fluor 647 goat anti-Mouse IgG1 (Cat# A-21240), Alexa Fluor 488 goat anti-Mouse IgG2b (Cat# A-21141: all from Thermo Fisher Scientific); or CyTM5 donkey anti-guinea pig (Cat# 706-175-148: Jackson Immunoresearch). The nuclei were counterstained with 4’,6-diamidino-2-phenylindole (DAPI: Thermo Fisher Scientific Cat D1306). The use of P2Y_13_, P2Y_1_ and ASCL1 antibodies required streptavidin-biotin amplification of the fluorescence signal. Finally, sections were mounted in PolySciences® mounting fluid.

For immunohistofluorescence (IHF) assays of the Ascl1 protein, brains were perfused with 25 mL of 2% PFA to avoid masking the epitope recognized by the antibody against Ascl1. The brains were equilibrated in 20% sucrose, and the sections obtained (25 μm) were permeabilized and blocked for 1 h at RT in IHF blocking solution: PBS, 10% normal goat serum (NGS), 1% Triton X-100 (v/v).

Fluorescence images of the tissue sections were taken on a Leica confocal microscope (model TCS SPE), controlled by Leica LAS AF software (version 2.5.1.5757), while fluorescence images of cells were obtained with a Nikon TE-2000 microscope controlled by Nikon’s NIS-Elements AR 4.5 software. Image processing was performed using ImageJ software.

### Fura-2 Microfluorimetry and Calcium Imaging

Calcium imaging experiments were carried out essentially as described previously (Carrasquero *et al*, 2005). Cells were seeded 24-48 hours earlier on 15 mm diameter glass coverslips and they were incubated for 45 min at 37 °C in Locke perfusion solution: 140 mM NaCl, 4.5 mM KCl, 2.5 mM CaCl_2_, 1.2 mM KH_2_PO_4_, 1.2 mM MgSO_4_, 5.5 mM glucose, 10 mM HEPES [pH 7.4], supplemented with 200 mg/mL BSA and 7.5 μM Fura-2 AM. The coverslips were then placed in a small superfusion chamber of a NIKON TE-200 microscope equipped with a Plan Fluor 20X/0.5 objective. The cells were continuously superfused with Locke’s medium at a rate of 1.5 mL/min using a hydrostatic pressure difference. Agonist stimulations were carried out for 30 seconds and pre-incubations with the antagonist for 5 minutes. Between stimulations the coverslips were washed with Locke perfusion for 5 min. The cells were illuminated alternately at wavelengths of 340 and 380 nm with a monochromator, which correspond to the excitation maxima of saturated and Ca^2+^-free acidic Fura-2 AM solutions, respectively. Fluorescence recordings were expressed as the ratio between the fluorescence values obtained at 340 and 380 nm as a function of time from an elliptical region above each cell analyzed, such that the F_340_/F_380_ ratio increases as the intracellular free Ca^2+^ concentration also increases. Before calculating the ratios, basal fluorescence values were quantified from a cell-free zone of the coverslip for each wavelength.

### Western blotting

SEZ tissue was dissected out in cold PBS, separating the dorsal from the ventral area. The tissue was lysed and then homogenized for 1-2 hours in a waterwheel at 4 °C in lysis buffer (pH 7.4) containing: 50 mM TrisHCl, 150 mM NaCl, 1% Nonidet P40 (all from Sigma-Aldrich), Complete™ Protease Inhibitor Cocktail Tablets (Roche Diagnostics GmbH) and 1 mM orthovanadate (Sigma-Aldrich). The resulting tissue lysate was centrifuged at 13,800 *g* for 15 minutes and the protein concentration was determined using a Bradford assay (Bio-Rad). The protein extracts were then separated by SDS-PAGE on 12% gels and transferred to polyvinylidene difluoride (PVDF) membranes, which were then blocked for 1h at RmT (25 °C) in a blocking solution of 3% BSA in TBS (10 mM Tris, 137 mM NaCl, 2.7 mM KCl, 5 mM Na_2_HPO_4_, 1.4 mM KH_2_PO_4_ and 0.1% Tween [pH 7.5]). The membranes were incubated with the primary antibodies raised against P2Y_13_ (1:100), P2Y_1_ (1:100) and α-Tubulin (1:10,000, Sigma-Aldrich Cat# T5168). Specific antibody binding was detected by ECL (enhanced chemiluminescence: Amersham GE Healthcare), and images were captured using ImageQuant LAS 500 (GE Healthcare Life Sciences) and analyzed with ImageQuant TL.

### Viral vector injections

Lentiviral constructs encoding P2Y_13_ and GFP were injected into both hemispheres of the SEZ of young adult (8-12 weeks) C57/B16 mice. The mice were anaesthetized (5 μg/mL Fentanyl, 0.5 mg/mL midazolam and 1 mg/mL medetomidine hydrochloride in saline) and injected with approximately 1 µL of the viral suspension at the stereotaxic coordinates relative to Bregma: 0.7 mm anteroposterior, 1.2 mm or −1.2 mm mediolateral and −1.6-2.1 mm dorsoventral. To awaken the mice, they were injected intraperitoneally (i.p., 10 μL/g mouse weight) with a solution of: 0.3 mg/mL buprenorphine, 0.25 mg/mL atipamezole hydrochloride and flumazenil 50 μg/mL in saline. The mice were then maintained individually in cages for 14 days with *ad libitum* access to food and water, in a temperature-controlled environment with 12-h light-dark cycles. After 14 days, histological sections of the brain were obtained.

Lenti-Cas9 virus was produced in HEK293T cells transfected with a mixture of the 3 transfection plasmids prepared and mixed with lipofectamine 2000 (Invitrogen) in a 1:3 ratio: packaging and enveloping plasmids psPAX2 (addgene#12260) and pMD2.VSVg (addgene#8454) together with lentiCas9-Blast plasmid (addgene#52962). After 72h, the medium was collected, centrifuged at 2000 rpm for 5 min and filtered with a 0.45 mm pore syringe filter.

Specific gRNAs targeting *P2RY13* (gRNA-P2ry13) sequence were selected based on the on-target and off-target scores provided by Benchling software (https://benchling.com/: San Francisco, CA, USA). The DNA sequences that produce gRNA targeting P2ry13 (P2ry13-1-F 5’-<u>aaac</u>ctcagacttgttgaagccctc-3’, P2ry13-1-R 5’-<u>caccg</u>agggcttcaacaagtctgag-3’) or a control sequence NTC (gNTC: NTC-F 5′-<u>caccg</u>cggctgaggcacctggttta-3′, NTC-R <u>aaac</u>taaaccaggtgcctcagccgc-3′) were then annealed and cloned into the BsmBI cloning site of lentiGuide-puro (Addgene #52963) plasmid that contains a U6 promoter driving the expression of the specific P2ry13 or NTC gRNA. Sanger sequencing confirmed the DNA insertion and plasmid sequence. Viral production to deliver gRNA to cells was performed as indicated above using the same packaging and enveloping plasmids together with lenti-Guide-puro containing gRNA-P2ry13 (gP2ry13) or control sequence. After 72h, the medium was collected, centrifuged at 2000 rpm for 5 min and filtered through a 0.45 mm pore syringe filter.

Specific CRISPR/Cas 9 activity using both virus at the *P2RY13* locus was tested in mouse cells lines and analyzed using ICE software (synthego) after PCR amplification followed by Sanger sequencing (Figure S6). After successfully testing, the subventricular area of mice (n=5) was inoculated with a total of 5 μl of the supernatant of a 1×10^7^ MOI/ml virus (ratio 1/10, Cas9/P2ry13 to ensure genetic edition).

### Whole-cell patch-clamp recordings

Whole-cell patch-clamp recordings were performed using an EPC10 amplifier and PatchMaster software (HEKA Electronic, Lambrecht, Germany). Patch micropipettes were pulled from borosilicate capillary tubes and adjusted to a final resistance of 4.5–5.5 MΩ once filled with the internal solution containing 145 mM KCl, 2 mM MgCl_2_, 0.3 mM EGTA, 0.3 mM GTP.Li_3_, and 10 mM HEPES (pH 7.2, ≈280 mOsm). The cells were immersed in an external solution composed of 145 mM NaCl, 2.8 mM KCl, 2 mM CaCl_2_, 1 mM MgCl_2_, 10 mM HEPES, and 10 mM glucose (pH 7.4, ≈300 mOsm), which was superfused continuously at approximately 2 mL/min through a recording chamber placed in the stage of a BX51W1 Olympus microscope.

Membrane currents were sampled at 10 kHz and filtered at 3 kHz, and series resistance manually by 80%. The p/n method with p/4 pulses was employed in all protocols involving rapid voltage changes for the automatic subtraction of capacitive and leak currents. The charge carried by voltage-activated currents was calculated as the time integral of the outward currents evoked by a voltage pulse to +10 mV (100 ms) from a holding potential (Vh) of −80 mV. Drugs were applied using a pneumatic ejection system (PDES-02DX, NPI Electronic GmbH, Germany) from a pipette with an opening of approximately 3–5 µm diameter positioned 5–10 µm from the cell under study. The experiments were carried out at RT (22–25 °C).

### Cell sorting

The SEZ from each hemisphere was dissected out as described previously (Belenguer et al., 2021) and each SEZ was cut into 5 or 6 small pieces of tissue and digested enzymatically using the Neural Tissue Dissociation Kit (T) in a GentleMACS Octo Dissociator with heating (Miltenyi Biotec). When the program was finished, 3 mL of trypsin inhibitor (10 mg/mL: Sigma) diluted in FACS buffer (calcium- and magnesium-free HBSS: 10 mM HEPES, 2 mM EDTA, 0.5% BSA) was added, and the tissue recovered by centrifugation. Subsequently, the mixture was pipetted up and down with a Pasteur pipette, between 20 and 25 times, until a homogeneous cell suspension was obtained. The samples were then filtered using a 40 μm strainer and centrifuged at 300g for 10 minutes. After discarding the supernatant, the pellet was resuspended with 30 μL of Myelin Removal Beads (Miltenyi) in 170 μL of FACS buffer, and incubated on ice for 15 minutes. The samples were then diluted in 1 mL of FACS buffer and centrifuged at 300g for 10 minutes. After resuspending with 0.5 mL, they were passed through a previously equilibrated LS column (Miltenyi) on a MidiMACS magnetic separator (Miltenyi). The columns were washed twice with FACS Buffer, and all the eluted fractions were collected and centrifuged again. The cell pellet was then incubated on ice for 30 minutes, protected from light, directly after resuspending in 100 μL of the antibody mix in FACS buffer: TER119-BV421 (BD, 1:200), CD31-BV421 (BD, 1:100), CD45-BV421 (BD, 1:100), and O4-405 (R&D, 1:50). Antibody binding was stopped by adding 1 mL of FACS buffer and centrifuging at 300g for 10 minutes., before resuspending each sample in 0.5 mL of FACS buffer. Finally, GFP^+^CD45^-^CD31^-^O4^-^Ter119^-^ cells were sorted on a BD FACSAria Fusion Cell Sorter (at 355 nm, 405 nm, 488 nm, 561 nm and 640 nm) and passed directly into 1.5 mL tubes containing 350 μL of RLT Plus for subsequent RNA extraction (Qiagen). Dead cells were excluded by staining with 0.1 ug/ml DAPI prior to analysis.

### FACS analysis

The SEZ of each animal was dissected out as described previously (Belenguer et al., 2021), cut into small pieces, and mechanically digested in 2 mL of FACS buffer by triturating in frosted glass pipettes until a homogeneous suspension was obtained. After filtering the samples through a 40 μm strainer, the cells were washed with FACS buffer (calcium- and magnesium-free HBSS, 10 mM HEPES, 2 mM EDTA, 0.5% BSA) using two consecutive rounds of centrifugation (300g, 10 min). The cell pellets recovered were resuspended in 100 μL of the antibody mix in FACS buffer for 30 minutes on ice, protected from light: CD45-BUV395 (1:100, BD), CD31-BV421 (1:100, BD), TER119-BV421 (1:200, BD), O4-AF405 (1:50, R&D), CD24-PerCP-Cy5.5 (1:300, BD), EGF-Alexa647 (1:300, Molecular Probes), GLAST-PE (1:50, Miltenyi), CD9-APC-Vio770 (1:20, Miltenyi), and P2Y13 (1:20, Alomone Labs). Antibody binding was stopped by adding 1 mL of FACS buffer, and the samples were centrifuged at 300g for 10 minutes. Finally, each sample was resuspended in 0.5 mL of FACS buffer. Dead cells were excluded by staining with DAPI (0.1 μg/mL) prior to the analysis, which was performed on a BD LSR-Fortessa cytometer with 355, 405, 488, 561 and 640 nm lasers.

### Statistical Analysis

The data obtained were represented as the arithmetic mean ± standard error of the mean (SEM) of independent experiments repeated a minimum of three times. The tests used were an analysis of variance ANOVA followed by Tukey’s method for the comparison of the means of the groups of values or of the means of groups with respect to the mean of one, respectively. An unpaired or paired Student’s t test was used to compare one variable between two groups. Statistical significance was established at a p-value <0.05 (*).

## ACKNOWLEDGMENTS

This work was supported by grants by Ministerio de Ciencia, Innovación y Universidades (MICIU, PID2022-138073OB-I00), and Red de Excelencia Consolider-Ingenio Spanish Ion Channel Initiative (BFU2015-70067REDC). The work described in this article was also supported by the COST Action CA21130 “P2X receptors as a therapeutic opportunity (PRESTO)”. AGU acknowledges funding from MICIU, (PID2020-117650RA-I00/ CNS2023-144109).

## AUTHOR CONTRIBUTIONS

Experiments were designed by FO. LP-H performed most of the experiments, analyzed the data and prepared the figures. JS-L, CLL-S and MLP-S contributed to the *in vivo* validation. DA-D contributed to the live imaging. LAO-O MA-B, ARA, and RG-V contributed to the functional analysis. AV-N and AR_P contributed to the bioinformatics analysis. AD-M,PD-A, and IF contributed to FACs and NSCs lineage progression analysis, DM-F and SG designed and tested the overexpression lentiviral constructs. PB and AG-U designed and tested the CRISPR/Cas9 lentiviral constructs. RP-S and EG.D contributed to P2Y_13_ expression analysis. FO drafted the manuscript.

## ACCESSION NUMBERS

ENA database under PRJEB77661 identifier

## DECLARATION OF CONFLICTS OF INTEREST

The authors declare no competing interests.

## Supplementary Figures

**Figure S1.**
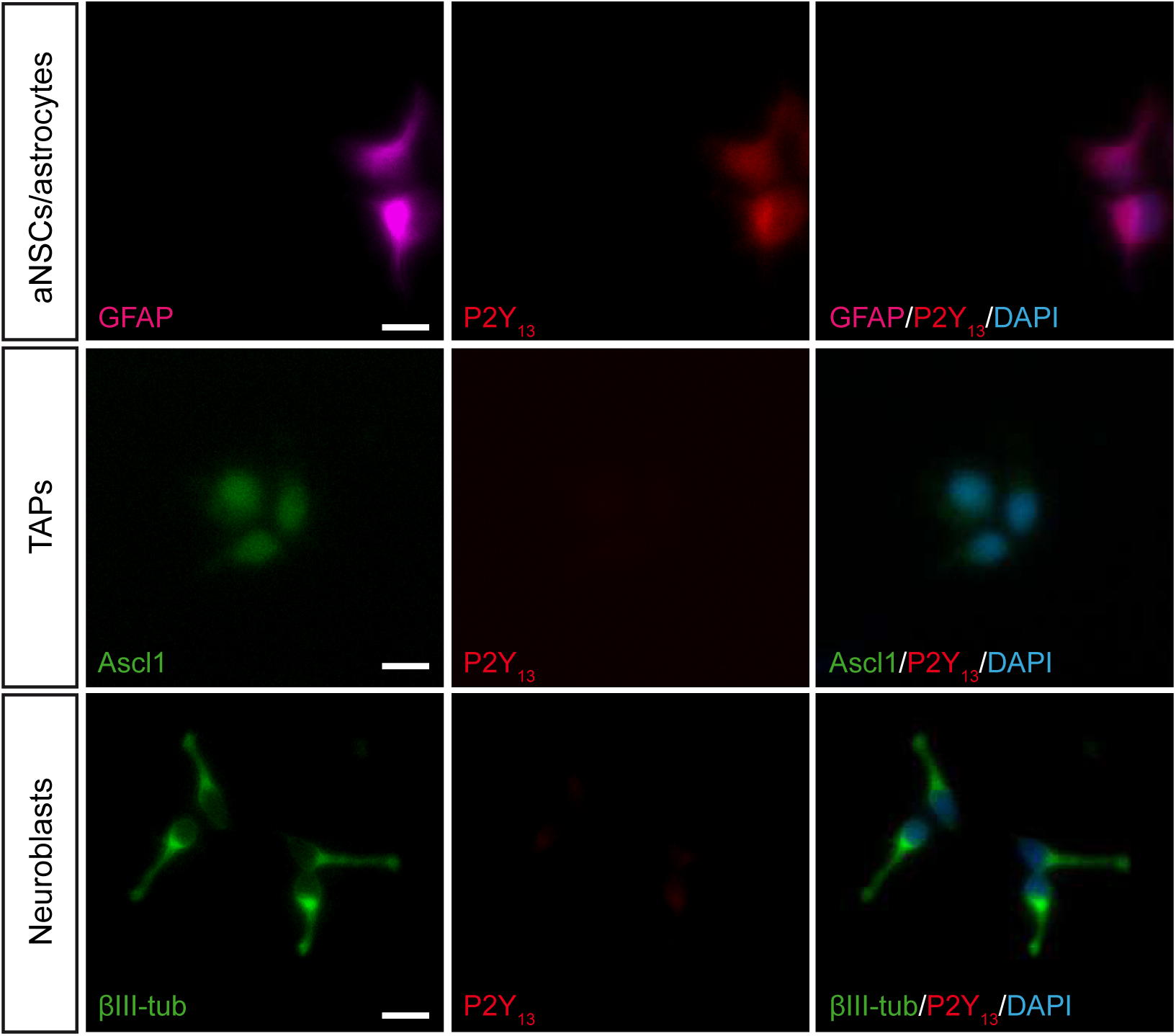
Expression of the P2Y_13_ receptor in the neurogenic lineage of the SEZ-derived cell cultures after 6 DIV: GFAP (magenta), Ascl1 (green, middle panel), βIII-tubulin (green, lower panel), P2Y_13_ (red). Note how P2Y_13_ receptor expression only co-localizes with GFAP in cells. Scale bar 30 µm.

**Figure S2.**
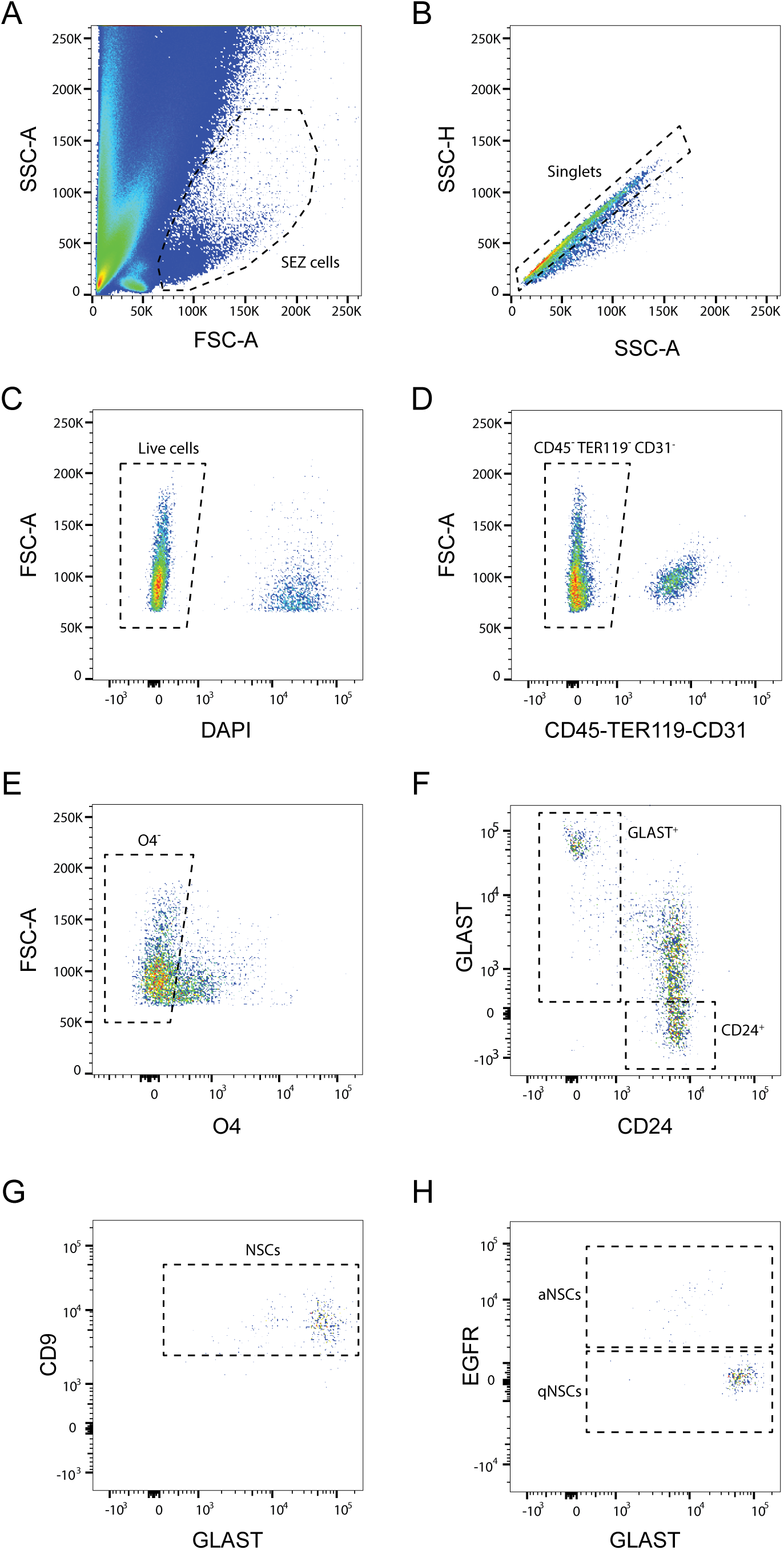
Flow cytometry gating strategy for the fractionation of the SEZ cell populations. **A.** FACS dot plot displaying forward scatter (FSC-A) versus side scatter (SSC-A) for the selection of the SEZ cells. **B.** FACS dot plot showing the side scatter-height (SSC-H) versus the side scatter-area (SSC-A) for doublet exclusion. **C.** FACS dot plot illustrating the DAPI staining used to exclude the dead cells. **D.** FACS dot plot presenting the intensity of CD45, CD31 and TER119 markers for the exclusion of endothelial cells, microglia and erythrocytes. **E.** FACS dot plot showing the intensity of O4 for the exclusion of oligodendrocytes. **F.** Representative FACS plot showing GLAST and CD24 staining in the Lin^-^ fraction. **G.** Within the GLAST^+^CD24^-^ fraction, CD9^high^ levels distinguish NSCs from non-neurogenic astrocytes. **H.** GLAST and EGFR levels define the GLAST^high^EGFR^-^ qNSCs and GLAST^low^EGFR^+^ aNSCs.

**Figure S3.**
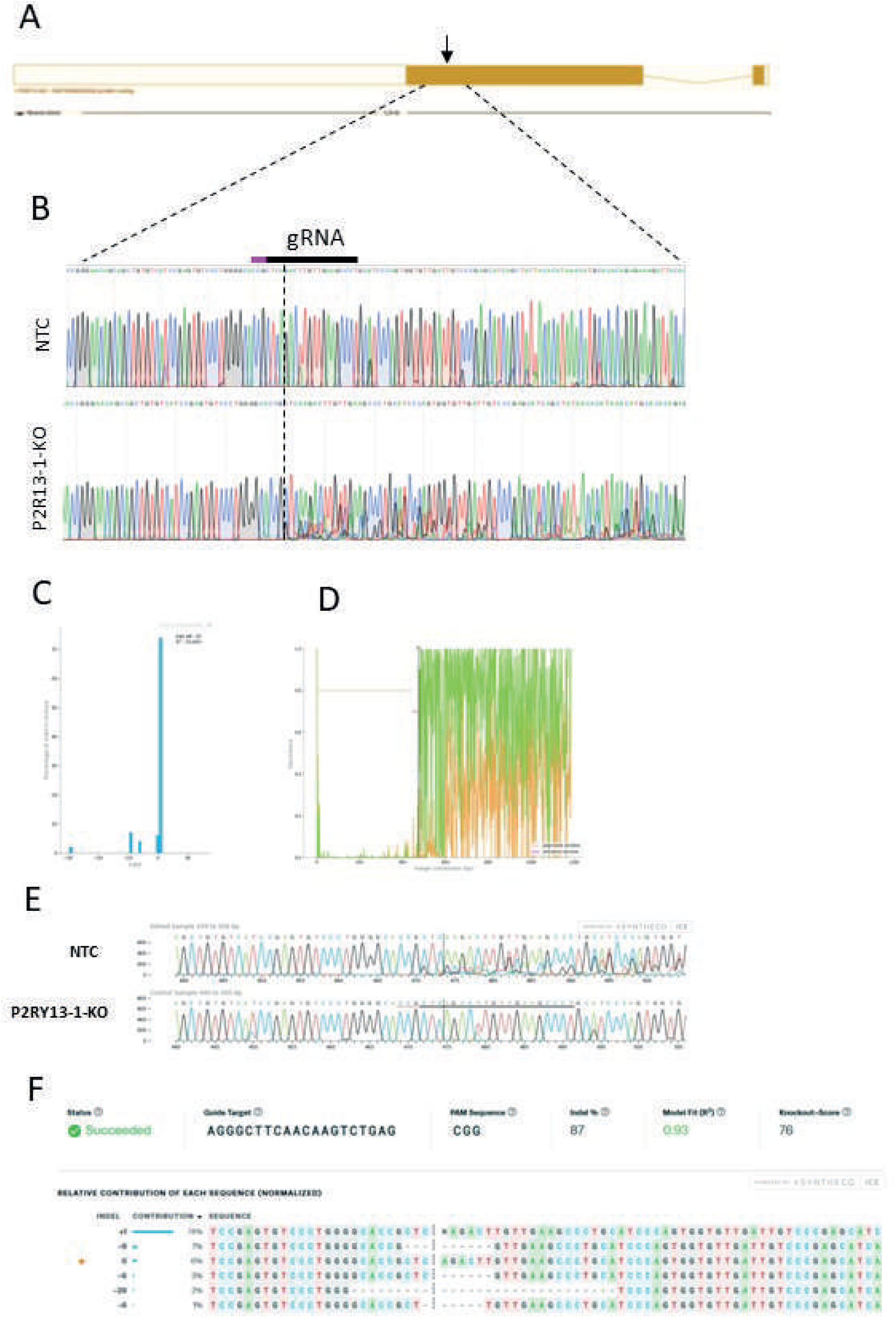
Sequence analysis of primary mouse cells infected with Cas9/sgRNA targeting the P2RY_13_ gene. **A.** Scheme of the targeted genome editing at the P2RY_13_ locus using CRISPR/Cas9. **B.** Sanger sequence traces of genomic DNA after the genomic edition in primary mouse cells. (**C-F**) **C-D.** The “Inference of CRISPR Edits” (ICE) software output analyzing the Sanger sequencing data from the P2RY_13_ gene showing the efficiency % (C), the Discordance and the Indel Distribution. **E.** alignments between the control and edited sequences across the first cut site. **F** the relative contribution of each sequence normalized.

**Figure S4.**
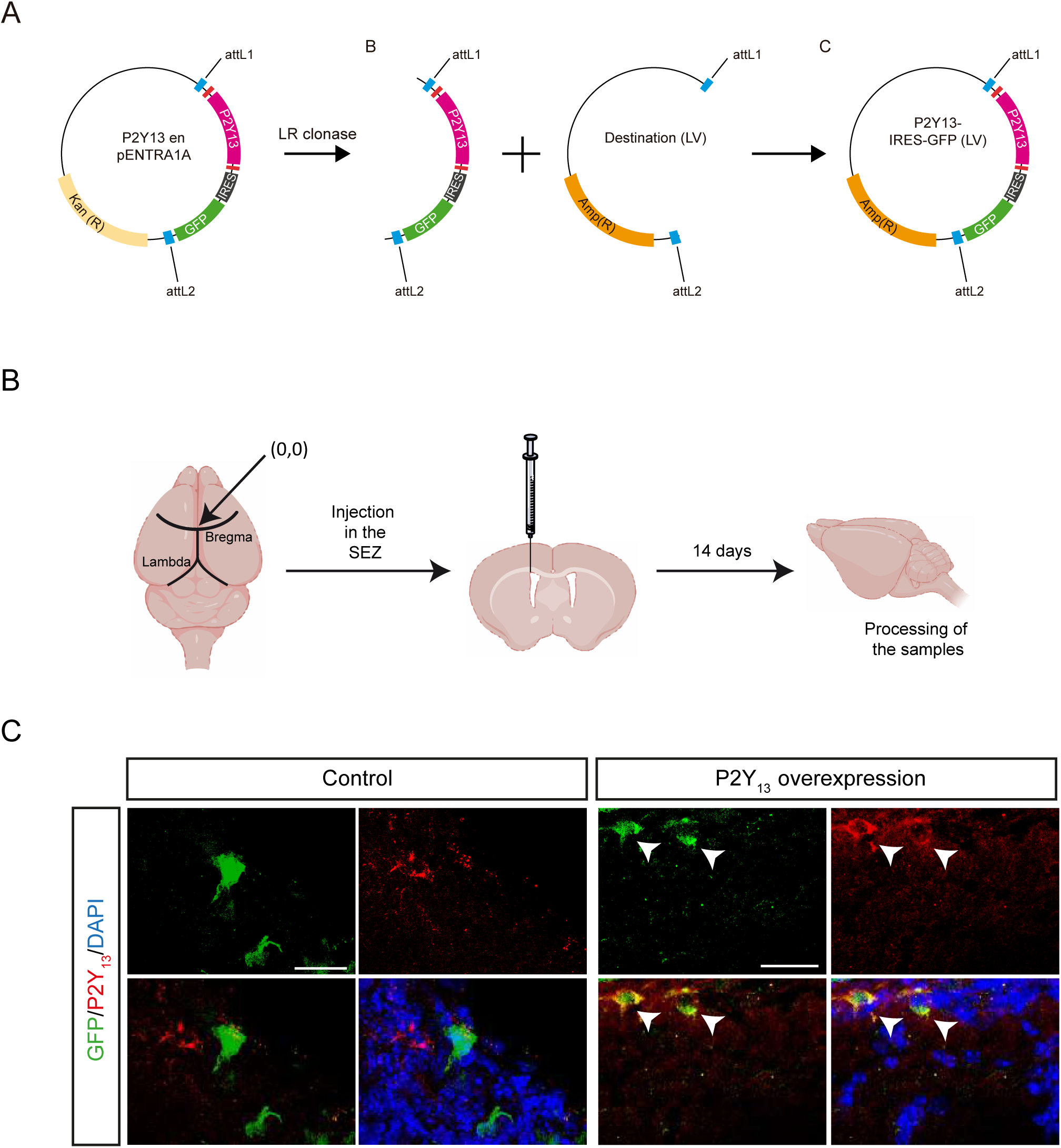
A. Scheme of the strategy to generate lentiviral vectors for the local overexpression of the P2Y_13_ receptor. **B**. Experimental design of lentiviral injection for local overexpression or silencing of the P2Y_13_ receptor. **C**. Positive control of the local overexpression of P2Y_13_ receptor. Lentiviral injection in the adult SEZ demonstrated that all LV-GFP-P2Y_13_ transduced cells (Green) co-localized with P2Y_13_ expression (red).

**Figure S5.**
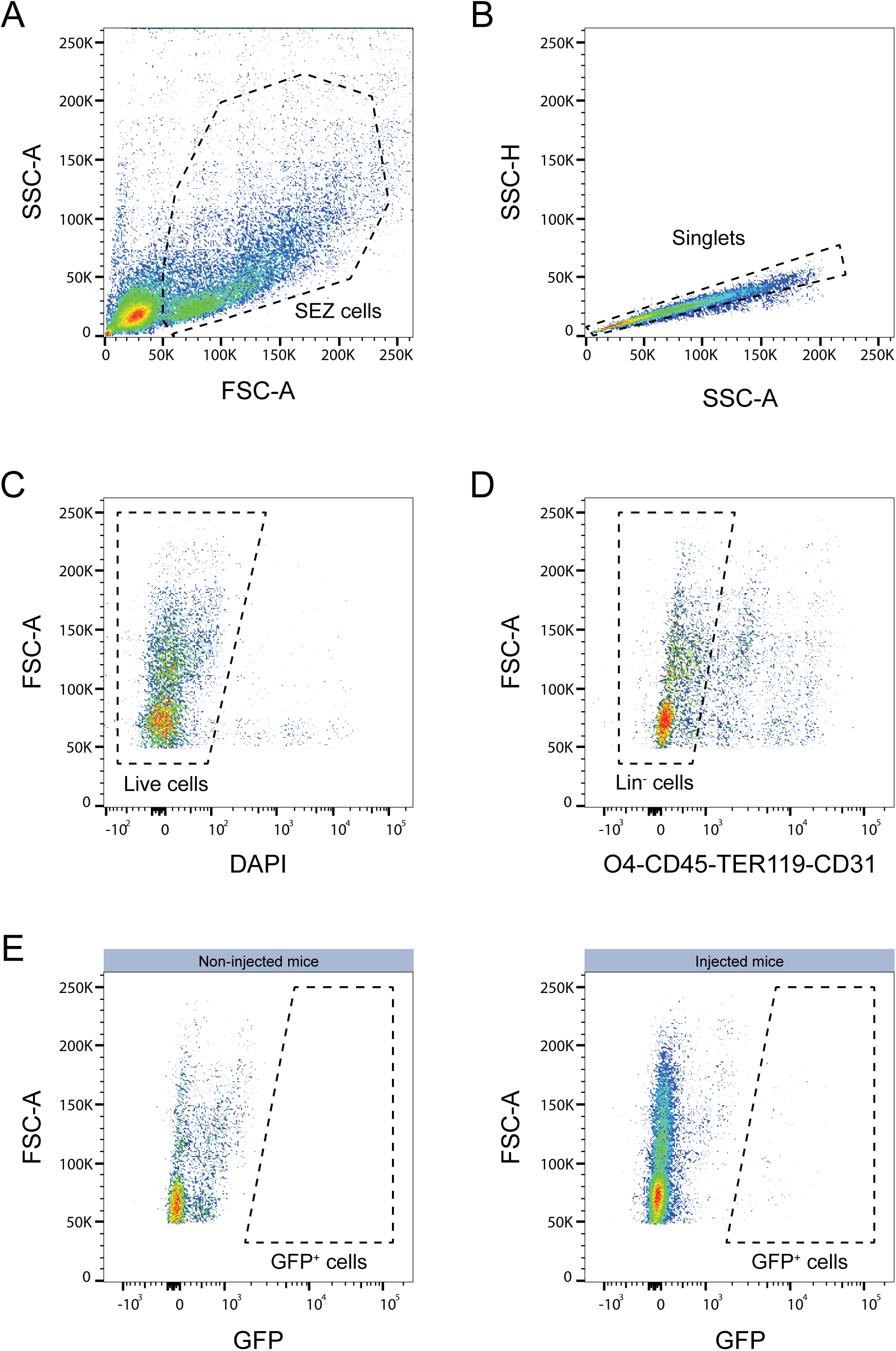
Flow cytometry gating strategy for GFP^+^Lin^-^ infected cell sorting. **A.** FACS dot plot displaying forward scatter (FSC-A) versus side scatter (SSC-A) for the selection of SEZ cells. **B.** FACS dot plot showing side scatter-height (SSC-H) versus side scatter-area (SSC-A) for doublet exclusion. **C.** FACS dot plot illustrating the DAPI staining used to exclude dead cells. **D.** FACS dot plot presenting the intensity of the CD45, CD31, TER119 and O4 markers for the exclusion of Lin⁺ cells. **E.** FACS dot plot comparing GFP intensity in non-injected mice (left) as a control versus lentivirus-injected mice (right).

**Figure S6.**
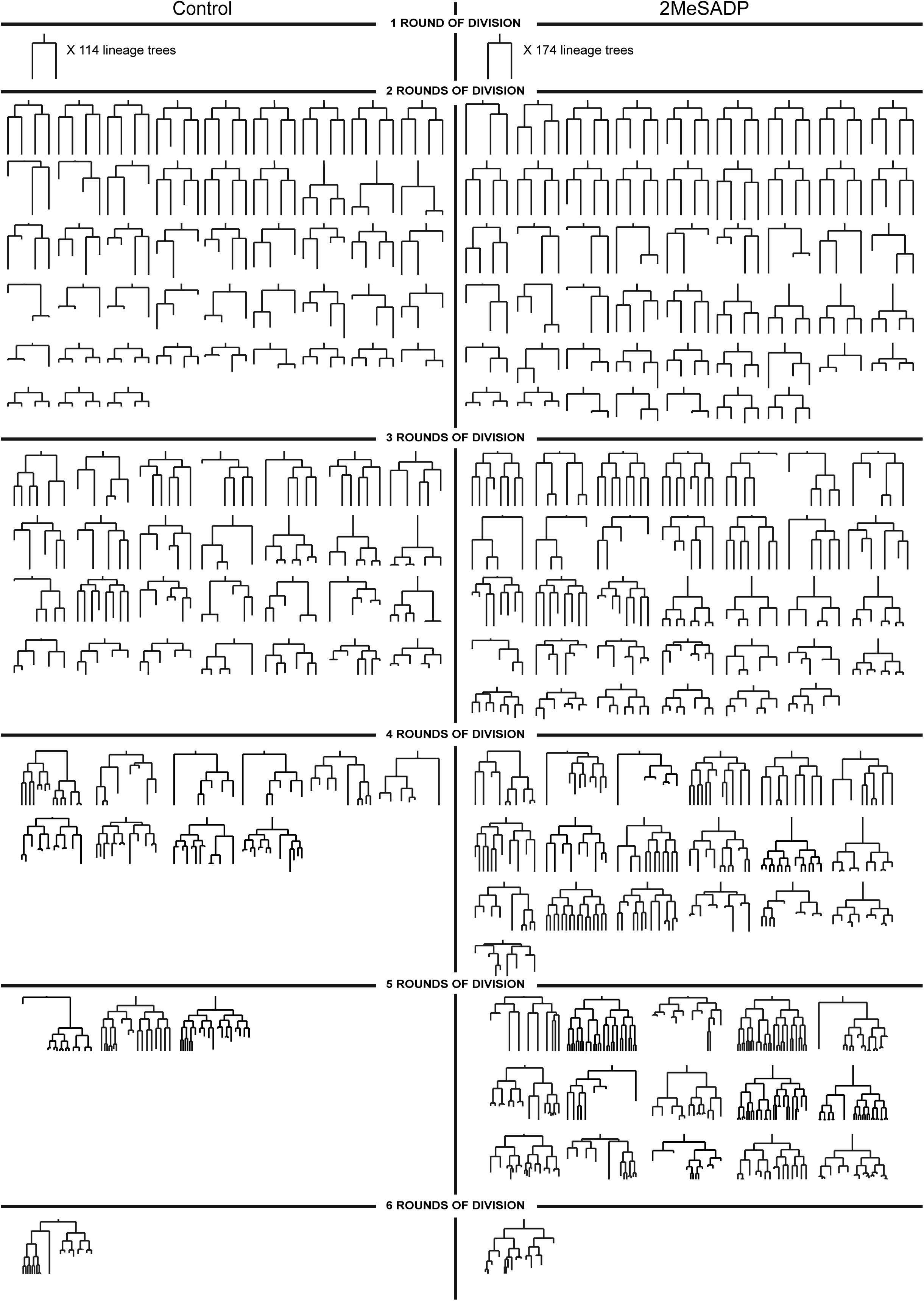
Summary of all the neurogenic clones tracked in control conditions or in 2MesADP-treated cultures. In each case, the identity of the progeny is determined by post-imaging immunocytochemistry.

**Figure S7.**
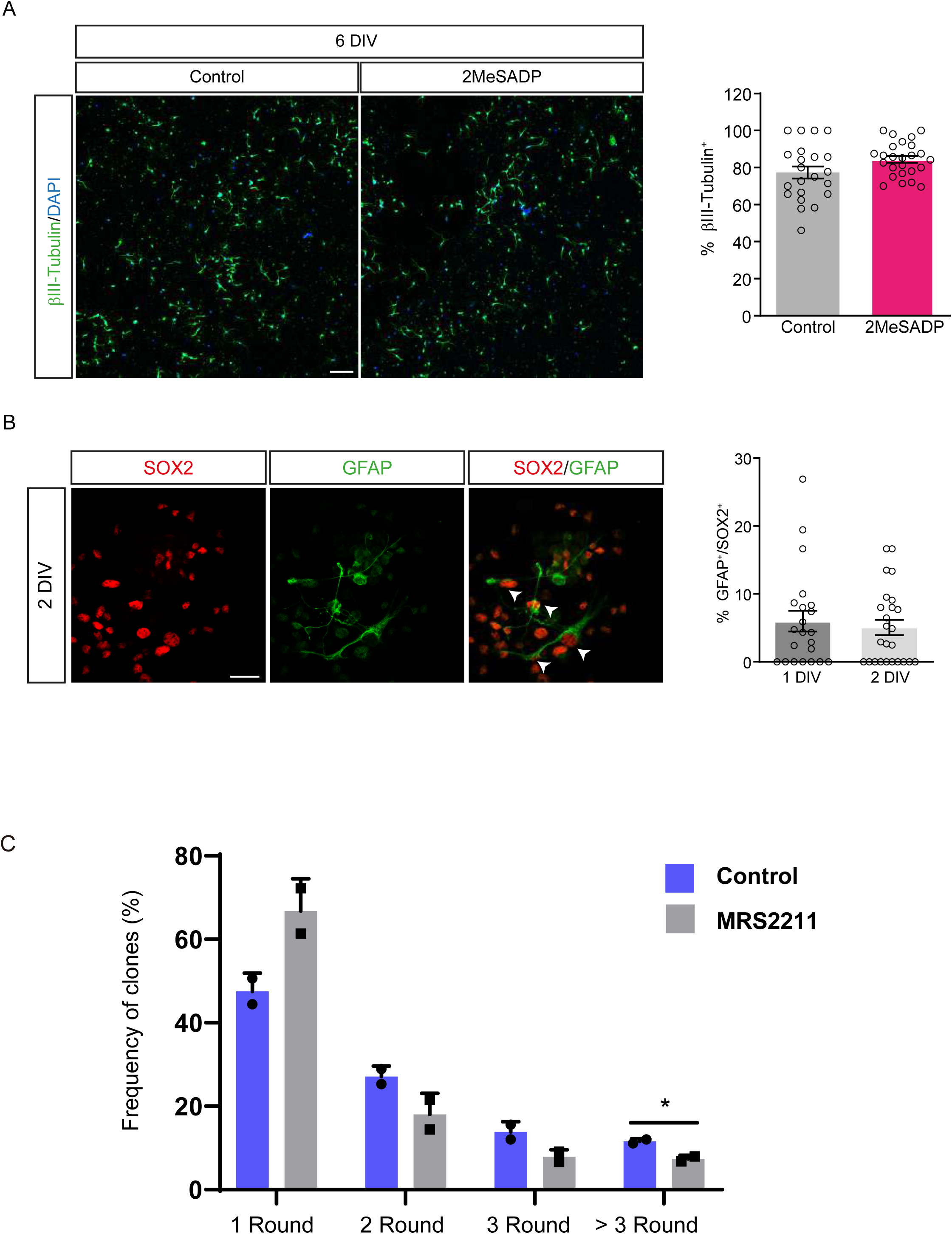
**A.** Quantification of the total number of βIII-tubulin^+^ (green) cells in SEZ cultures after 6 DIV, in either control conditions or in the presence of 2MeSADP (n=3). Scale bar 100 µm. **B.** Quantification of the dual labelled GFAP^+^ (Green) and SOX2^+^ (red) cells in SEZ cultures after 1 and 2 DIV (n=2). Scale bar 30 µm. All graphs show the mean ±SEM. **C.** Percentage of clones undergoing 1-3 or >3 rounds of division in the live imaging experiments under control conditions or in the presence of MRS2211 (n=2). The graphs show the mean ±SEM: *p<0.05 (T-test).

## Notes

### Competing Interest Statement

The authors have declared no competing interest.

### Summary of Updates

Although the final version is still under revision, we have included Lucia Gallego in this preprint to acknowledge her contribution to specific aspects of the work.

